# Dynamic reconfiguration of brain coactivation states associated with active and lecture-based learning of university physics

**DOI:** 10.1101/2025.02.22.639361

**Authors:** Donisha D. Smith, Jessica E. Bartley, Julio A. Peraza, Katherine L. Bottenhorn, Jason S. Nomi, Lucina Q. Uddin, Michael C. Riedel, Taylor Salo, Robert W. Laird, Shannon M. Pruden, Matthew T. Sutherland, Eric Brewe, Angela R. Laird

**Author notes:** Corresponding Author*: **Donisha D. Smith,** Department of Psychology, Florida International University Miami, FL, USA.

## Abstract

Academic institutions are increasingly adopting active learning methods to enhance educational outcomes. Using functional magnetic resonance imaging (fMRI), we investigated neurobiological differences between active learning and traditional lecture-based approaches in university physics education. Undergraduate students enrolled in an introductory physics course underwent an fMRI session before and after a 15-week semester. Coactivation pattern (CAP) analysis was used to examine the temporal dynamics of brain states across different cognitive contexts, including physics conceptual reasoning, physics knowledge retrieval, and rest. CAP results identified seven distinct brain states, with contributions from frontoparietal, somatomotor, and visuospatial networks. Among active learning students, physics learning was associated with increased engagement of a somatomotor network, supporting an embodied cognition framework, while lecture-based students demonstrated stronger engagement of a visuospatial network, consistent with observational learning. These findings suggest significant neural restructuring over a semester of physics learning, with different instructional approaches preferentially modulating distinct patterns of brain dynamics.

## Introduction

Physics education research has led to the development of evidence-based instructional strategies aimed at effectively disseminating physics concepts to students, fostering deeper conceptual understanding, minimizing route-learning and non-divergent thinking, and enabling the construction of expert-like knowledge structures that enable students to apply physics principles to real-world problems ^1,2^. Among these methodologies, active learning has emerged as a pivotal strategy for developing a more integrative understanding of physics among students and has seen widespread adoption throughout various academic institutions^3^. Active learning strategies adopt a student-centered approach, encouraging students to engage in group activities and collaboratively teach one another relevant course material with minimal instructor intervention^4,5^. In contrast, traditional, passive learning environments are characterized by an instructor-centered model, wherein the educator delivers course content to students, who are expected to process the material through listening, note-taking, and occasional participation during lectures at the instructor’s discretion^4,6^.

Despite the numerous behavioral studies assessing the differences between active and passive learning methodologies in physics education^7–11^, the longitudinal neurocognitive effects of these different instructional methods remain largely unexplored. These fundamentally different instructional strategies may selectively modulate distinct functional brain networks, which in turn could affect the underlying neural representations that are engaged when students interact with and internalize physics-related content and complex physics concepts. In contrast to passive learning, active learning emphasizes collaborative, hands-on engagement with learning materials, typically leading to enhanced conceptual understanding ^12–14^. Consequently, a theoretical framework that may provide insights into the anticipated differences in the neurobiological mechanisms of active versus passive physics learning is embodied cognition, which posits that cognitive processes are deeply rooted in the body’s interactions with the world^15,16^. According to this framework, conceptual knowledge is grounded in sensory and motor experiences, and understanding abstract concepts involves simulating sensorimotor patterns associated with those concepts^17^.

Prior neuroimaging research has shown that processing action-related words and concepts activates sensorimotor regions of the brain, a phenomenon known as semantic somatotopy ^18–20^. This suggests that engaging in physical experiences can enhance the comprehension of abstract concepts by recruiting sensorimotor neural networks. In the context of physics education, students who actively and physically interact with learning materials may develop stronger and more integrative neural activation patterns when engaging with physics concepts. For instance, Kontra et al.^19^ found that university students who learned about angular momentum through interactive physical models exhibited greater activation in brain regions associated with action planning and motor experiences, such as the dorsal premotor cortex, primary motor/somatosensory cortex, superior parietal lobe, and supplementary motor area. Moreover, the involvement of sensorimotor regions in understanding abstract physics concepts may result from the neural reuse of regions initially dedicated to lower-level sensory and motor functions now supporting higher-order cognitive processes^21^. This neural reuse is evident in studies demonstrating that regions in the parietal and premotor cortex are implicated in mental rotation tasks and spatial reasoning, both critical cognitive domains in physics learning, engaging brain networks such as the somatomotor, dorsal attention, and frontoparietal networks^22^.

Evidence supporting embodied cognition in physics learning suggests that different teaching methods may create distinct patterns of brain network activations over time. Prior neuroimaging evidence has shown that learning involves dynamic brain network reconfiguration^23^. Methods that capture time-varying properties of neuroimaging data are known to align more strongly with shifts in cognition and behavior than static functional connectivity (sFC) techniques^24^. Thus, these methods may provide more informative insights on the neurobiological differences between active and passive instruction. SFC techniques analyze functional connectivity within and between brain regions while assuming that the observed connectivity patterns remain stationary throughout the entire data acquisition period^24,25^. This approach overlooks the dynamic nature of brain connectivity, which often demonstrates considerable temporal variability within seconds during different cognitive tasks and learning processes^26^. Capturing these temporal dynamics is particularly important as changes in functional connectivity over time may reflect underlying neuroplastic processes such as Hebbian learning, which posits that persistent activation of specific neural patterns in response to repeated engagement with tasks leads to intrinsic cortical restructuring and stable, recurrent states^27–29^. Moreover, prior research on metastable brain states, defined as transient patterns of network activations that are observed at greater frequencies during resting state, suggests that higher metastability of certain brain states during rest is associated with increased pre-configuration towards tasks that rely on the underlying cognitive processes related to those neural patterns^30–32^. Hence, by exploring temporal dynamics through dynamic functional connectivity (dFC) methods, transient patterns of network activations can be captured, identifying potential neural restructuring associated with different instructional methods over time and providing greater insights that may be overlooked by traditional static methods.

To capture these dynamic patterns, we leveraged the coactivation pattern (CAP) analytical technique, a dynamic method that aggregates similar spatial distributions of brain activity^33,34^ to examine the temporal dynamics of brain networks across different cognitive contexts, including physics conceptual reasoning, physics knowledge retrieval, and rest. Brain states consistently identified across multiple cognitive contexts are likely to provide a robust representation of intrinsic functional connectivity that is more generalizable, reliable, and meaningful compared to states identified from single task fMRI, especially when broad networks (e.g., default, frontoparietal, somatomotor, and attention networks) are of interest^35–37^. We utilized three reproducible CAP metrics^38^: 1) temporal fraction, defined as the proportion of time occupied by a single state; 2) persistence, defined as the average duration a state persists before transitioning to another state; and 3) counts, defined as the frequency of CAP initiation (contiguous CAPs are considered a single initiation) across the entire scan.

In the present study, we investigated the neurobiological differences in dFC among students who completed a semester of an introductory undergraduate physics course in either active learning or traditional lecture-based classrooms. To this end, we assessed different cognitive contexts, using two in-scanner fMRI tasks, including the Force Concept Inventory (FCI), a physics-related conceptual reasoning task examining motion trajectories of objects at rest or in motion and the Physics Knowledge (PK) task, a physics-related semantic retrieval task assessing general physics knowledge, as well as resting state. Across these different contexts, we examined how time (pre- versus post-instruction) and instructional approach (active learning versus lecture-based classrooms) were associated with different brain network activation patterns. After a semester of physics instruction, we expected students to show increased engagement of brain regions critical for physics learning, particularly the frontoparietal, somatomotor, and visual networks that support spatial comprehension and mathematical thinking^39,40^. However, we anticipated that the method of instruction would preferentially modulate distinct neural patterns. Drawing on embodied cognition theory and principles of Hebbian learning, we hypothesized that active learning students would develop more integrated network activation patterns, evidenced by greater temporal fraction, persistence, and counts in CAPs characterized by widespread activation across multiple networks, including increased somatomotor engagement. We anticipated that this multimodal pattern would reflect how active learning facilitates comprehension of physics understanding through collaborative physical interaction with concepts. In contrast, we expected lecture-based students to show increased temporal fraction, persistence, and counts in CAPs dominated by visual network activation, reflecting how these students primarily acquire physics knowledge through observation ^41^. Finally, we predicted that instructional method-specific patterns would be most observed during resting state, suggesting that different teaching approaches are linked to intrinsic reconfiguration that generalizes beyond physics-related cognition during domain-specific tasks.

## Results

### Demographic Differences Between Active Learning and Lecture-Based Classrooms

The study sample comprised 121 undergraduate students enrolled in a 15-week introductory physics course that employed either a modeling instruction approach (n=61), emphasizing model building and collaborative group activities^42^, or a traditional lecture-based instruction approach (n=60). Behavioral and neuroimaging data were collected at two time periods: pre- and post-instruction. We assessed for potential demographic differences between students in the active learning classrooms (*n*=61) and those in the lecture-based classrooms (*n*=60) and found no significant differences in age, sex, ethnicity, household income, grade point average (GPA), and years enrolled at FIU (i.e., freshman, sophomore, junior, or senior) (**Supplementary Table 2**). We further examined a subsample of participants with at least one functional run for all tasks and sessions, which included active learning (*n*=46) and lecture-based students (*n*=44), and also observed no significant demographic differences (**Supplementary Table 2**).

### Spatial Topography of Coactivation Patterns (CAPs)

We identified seven CAPs that dynamically varied at both pre- and post-instruction during the task and rest runs (**Fig. 1**); this number of brain states was selected using the elbow criterion (**Supplementary Fig. 1**). These patterns were identified through k-means clustering of concatenated resting state and task data (FCI and PK) from all participants with at least one functional run across all tasks and sessions (n=90). The spatial dimensionality of the extracted time series data from each participant was reduced using the HCPex atlas ^43^, an extended MNI version of the Glasser et al. ^44^ parcellation, which includes 66 subcortical regions, resulting in 426 nodes across 22 cortical divisions. “High Amplitude” cosine similarities quantify the similarity between a cortical division from the HCPex atlas and the positive activations within a CAP cluster centroid. These values represent regions that are more active than average. Conversely, “Low Amplitude” cosine similarities measure the similarity between a cortical division and the negative activations within the cluster centroid, indicating regions with activity levels below the mean. While the “Low Amplitude” cosine similarity are negative values, for visualization purposes, the absolute values of the negative activations from each CAP cluster centroids were computed to restrict all cosine similarity values from 0 to 1. However, the negative cosine similarities are reported below to differentiate them from the cosine similarities of the “High Amplitude” results. Finally, we then characterized each CAP by the network correspondence between HCPex cortical divisions exhibiting a cosine similarity ≥ 0. 20 to the positive activations in the CAP and 12 resting state networks from the Cole-Anticevic atlas^45^ through spin permutation^46,47^.

**Figure 1.**
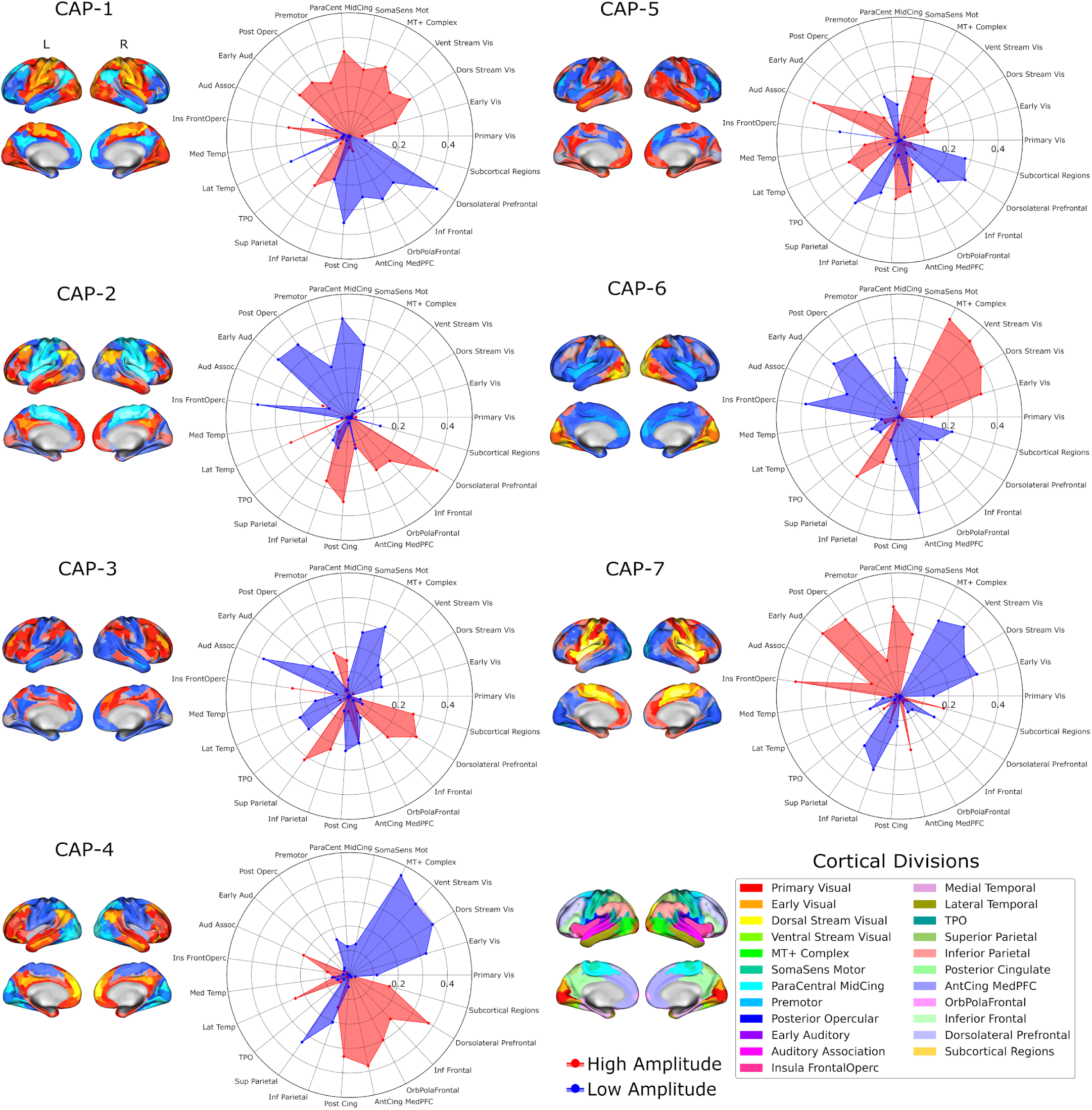
Cosine Similarity Analysis of CAPS. Surface plots for each CAP are paired with radar plots depicting the cosine similarity between each CAP and various HCPex cortical divisions. In these radar plots, “High Amplitude” (shown in red) represent the cosine similarity between the cortical division and the positive activations in the cluster centroid, while “Low Amplitude” (shown in blue) represent the cosine similarity between the cortical division and the negative activations in a CAP cluster centroid. The radial axis of each radar plot shows cosine similarity values ranging from 0 to 0.5. Note that during computation, the inverse values of the negative activations were calculated to restrict the cosine similarity for both positive and negative activations in a CAP from 0 to 1. At the bottom right, a depiction of each HCPex cortical division (excluding the subcortical areas) is shown. The labels for each CAP show the network correspondence of cosine similarities ≥ 0. 20.

**CAP-1** showed widespread activations across the Paracentral Midcingulate (*cosine similarity* [*cos*(θ)] = 0. 34), MT+ Complex (*cos*(θ) = 0. 31), Dorsal Stream Visual (*cos*(θ) = 0. 28), Somatosensory regions (*cos*(θ) = 0. 27), Posterior Opercular (*cos*(θ) = 0. 26), Early Auditory (*cos*(θ) = 0. 26), Insula Frontal Opercular (*cos*(θ) = 0. 25), Superior Parietal (*cos*(θ) = 0. 25), Ventral Stream Visual (*cos*(θ) = 0. 24), and the Premotor (*cos*(θ) = 0. 23). These patterns demonstrated significant correspondence with the **Somatomotor Network** (*Dice Similarity Coefficient* [*DSC*] = 0. 4875, *p* = 0. 001) and the **Cingulo-Opercular Network** (*DSC* = 0. 3006, *p* = 0. 023). Dorsolateral Prefrontal Cortex (*cos*(θ) = − 0. 41), Posterior Cingulate (*cos*(θ) = − 0. 35), OrbPolaFrontal (*cos*(θ) = − 0. 29), Lateral Temporal (*cos*(θ) = − 0. 26), Inferior Frontal regions (*cos*(θ) = − 0. 26), and the Anterior Cingulate Medial Prefrontal Cortex (*cos*(θ) = − 0. 25).

**CAP-2** demonstrated robust activations in the Dorsolateral Prefrontal Cortex (*cos*(θ) = 0. 42), Posterior Cingulate (*cos*(θ) = 0. 34), Inferior Parietal (*cos*(θ) = 0. 27, Lateral Temporal (*cos*(θ) = 0. 26), Inferior Frontal (*cos*(θ) = 0. 24), and the Orbitofrontal Cortex (*cos*(θ) = 0. 24). These activation patterns showed strong correspondence with both the **Default Mode Network** (*DSC* = 0. 4898, *p* = 0. 001) and the **Frontoparietal Network** (*DSC* = 0. 4511, *p* = 0. 001). Notable deactivations were observed in the Paracentral Midcingulate (*cos*(θ) = − 0. 40), Insula Frontal Opercular (*cos*(θ) = − 0. 37), Early Auditory (*cos*(θ) = − 0. 37), Posterior Opercular (*cos*(θ) = − 0. 36), Somatosensory Motor (*cos*(θ) = − 0. 30), and the Premotor (*cos*(θ) = − 0. 22).

**CAP-3** exhibited significant activations in the Dorsolateral Prefrontal Cortex (*cos*(θ) = 0. 32), Superior Parietal (*cos*(θ) = 0. 31), Subcortical (*cos*(θ) = 0. 27), Insula Frontal Opercular (*cos*(θ) = 0. 23), Inferior Parietal (*cos*(θ) = 0. 23), and the Inferior Frontal (*cos*(θ) = 0. 23). These activation patterns aligned significantly with the **Frontoparietal Network** (*DSC* = 0. 3940, *p* = 0. 001), **Cingulo-Opercular Network** (*DSC* = 0. 3287, *p* = 0. 013), and the **Dorsal Attention Network** (*DSC* = 0. 1629, *p* = 0. 007). Substantial deactivations were observed in the Auditory Association regions (*cos*(θ) = − 0. 37), MT+ Complex (*cos*(θ) = − 0. 32), Somatosensory (*cos*(θ) = − 0. 27), Posterior Cingulate (*cos*(θ) = − 0. 22), Lateral Temporal (*cos*(θ) = − 0. 22), and the Temporoparietal-Occipital junction (*cos*(θ) = − 0. 21).

**CAP-4** exhibited pronounced activations in the Anterior Cingulate Medial Prefrontal Cortex (*cos*(θ) = 0. 38), Dorsolateral Prefrontal Cortex (*cos*(θ) = 0. 38), and Posterior Cingulate (*cos*(θ) = 0. 33), alongside the OrbPolaFrontal (*cos*(θ) = 0. 30), Inferior Frontal (*cos*(θ) = 0. 24), Lateral Temporal (*cos*(θ) = 0. 24), and the Auditory Association (*cos*(θ) = 0. 20). These patterns showed significant correspondence with the **Default Mode Network** (*DSC* = 0. 5440, *p* = 0. 001), **Frontoparietal Network** (*DSC* = 0. 3647, *p* = 0. 001), and the **Language Network** (*DSC* = 0. 1726, *p* = 0. 015). Strong deactivations were noted across visual processing areas, particularly in the MT+ Complex (*cos*(θ) = − 0. 46), Dorsal Stream Visual (*cos*(θ) = − 0. 40), Ventral Stream Visual (*cos*(θ) = − 0. 40), Superior Parietal (*cos*(θ) = − 0. 33), Early Visual (*cos*(θ) = − 0. 32), and the Inferior Parietal (*cos*(θ) = − 0. 20).

**CAP-5** demonstrated robust activations in the Auditory Association (*cos*(θ) = 0. 38), MT+ Complex (*cos*(θ) = 0. 28), Somatosensory Motor (*cos*(θ) = 0. 26), Posterior Cingulate (*cos*(θ) = 0. 24), Lateral Temporal (*cos*(θ) = 0. 22), and the Anterior Cingulate Cortex (*cos*(θ) =− 0. 21). These activation patterns aligned significantly with the **Default Mode Network** (*DSC* = 0. 4316, *p* = 0. 004) and **Somatomotor Network** (*DSC* = 0. 2712, *p* = 0. 002). Notable deactivations were observed in the Superior Parietal (*cos*(θ) = − 0. 31), Dorsolateral Prefrontal Cortex (*cos*(θ) = − 0. 31), Subcortical (*cos*(θ) = − 0. 27), Insula Frontal Opercular (*cos*(θ) = − 0. 24), Inferior Frontal (*cos*(θ) = − 0. 23), and the Inferior Parietal (*cos*(θ) = − 0. 23).

**CAP-6** revealed extensive activations across visual processing regions, including MT+ Complex (*cos*(θ) = 0. 45), Ventral Stream Visual (*cos*(θ) = 0. 42), Dorsal Stream Visual (*cos*(θ) = 0. 39), Early Visual (*cos*(θ) = 0. 34), and the Superior Parietal (*cos*(θ) = 0. 30). These activation patterns showed strong correspondence with the **Secondary Visual Network** (*DSC* = 0. 7102, *p* = 0. 001) and the **Dorsal Attention Network** (*DSC* = 0. 1406, *p* = 0. 036). Pronounced deactivations were observed in Anterior Cingulate Medial Prefrontal Cortex (*cos*(θ) = − 0. 40), Insula Frontal Opercular (*cos*(θ) = − 0. 38), Early Auditory (*cos*(θ) = − 0. 34), Posterior Opercular (*cos*(θ) = − 0. 31), Paracentral Midcingulate (*cos*(θ) = − 0. 24), Auditory Association (*cos*(θ) = − 0. 23), and the Subcortical (*cos*(θ) =− 0. 22).

**CAP-7** revealed strong activations in Insula Frontal Opercular (*cos*(θ) = 0. 42), Early Auditory (*cos*(θ) = 0. 40), Posterior Opercular (*cos*(θ) = 0. 38), Paracentral Midcingulate (*cos*(θ) = 0. 36), Somatosensory Motor (*cos*(θ) = 0. 25), and the Anterior Cingulate Medial Prefrontal Cortex (*cos*(θ) = 0. 22). These patterns demonstrated significant overlap with the **Somatomotor Network** (*DSC* = 0. 4679, *p* = 0. 001) and the **Cingulo-Opercular Network** (*DSC* = 0. 3517, *p* = 0. 005). Prominent deactivations were observed in the Ventral Stream Visual (*cos*(θ) = − 0. 38), MT+ Complex (*cos*(θ) = − 0. 34), Early Visual (*cos*(θ) = − 0. 32), Inferior Parietal (*cos*(θ) = − 0. 32), Dorsal Stream Visual (*cos*(θ) = − 0. 31), and the Superior Parietal (*cos*(θ) = − 0. 25).

Comparison of the spatial patterns for the seven CAPs revealed complex relationships among them (**Fig. 2**). Notably, each CAP demonstrated significant anti-correlation with at least one other CAP. CAP-1 showed significant negative correlations with CAP-2 and CAP-4. CAP-2 was found to be strongly anti-correlated with CAP-7. CAP-3 had a significant negative correlation with CAP-5, while CAP-4 was anti-correlated with CAP-6 and CAP-7. Additionally, CAP-6 and CAP-7 exhibited a notable anti-correlation with each other.

**Figure 2.**
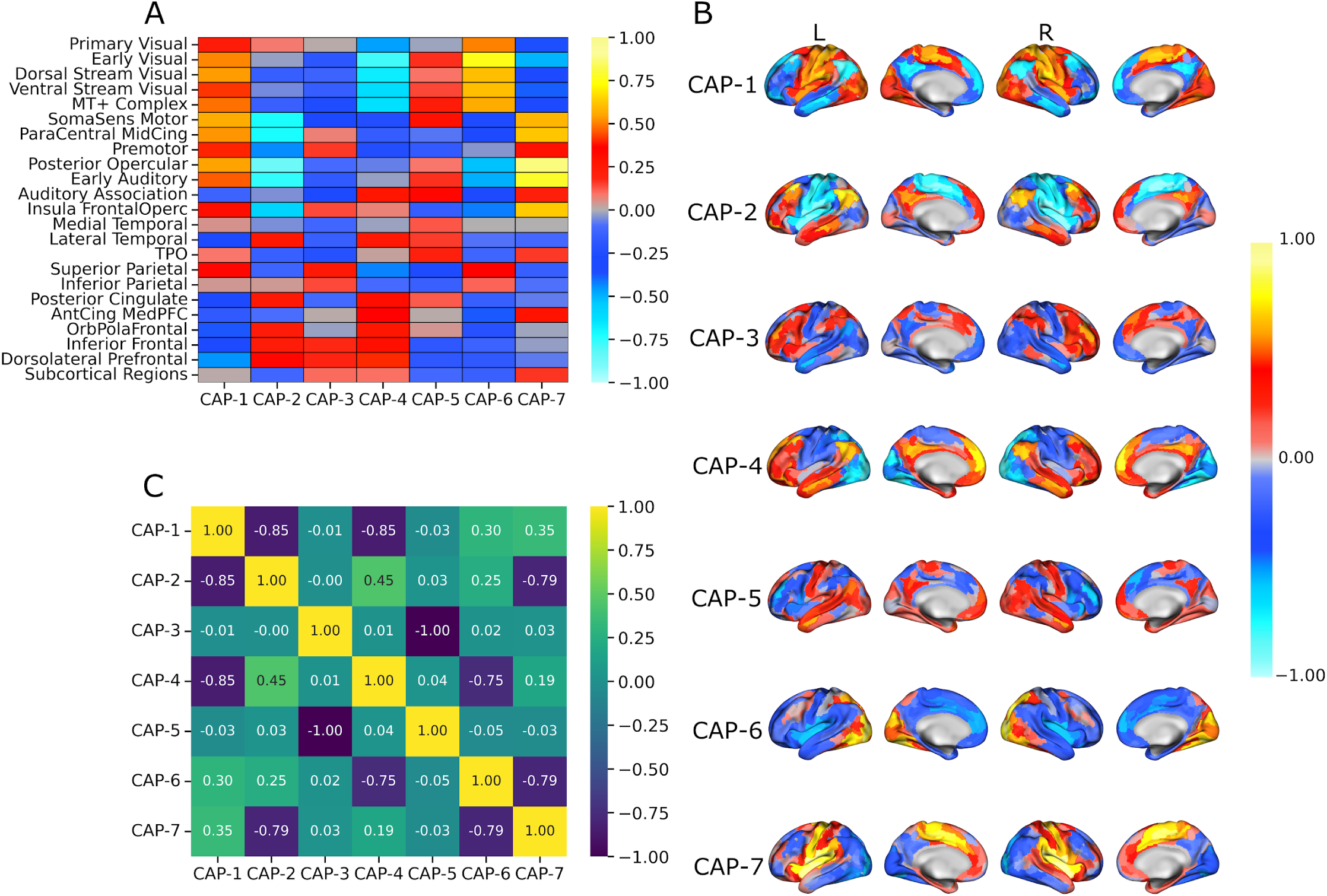
Visualizations of Coactivation Patterns. (A) A heatmap depicts the mean activation of HCPex cortical divisions across seven identified CAPs. Red indicates higher activation, while blue represents lower activation or deactivation. (B) Surface plots illustrate the spatial distribution of activation (yellow) and deactivation (blue) for each CAP across left (L) and right (R) hemispheres. (C) A correlation matrix shows the Pearson correlation coefficients between all pairs of CAPs. The color scale ranges from purple (negative correlation) to yellow (positive correlation). Notably, each CAP demonstrated significant anti-correlation with at least one other CAP. CAP-1 showed significant negative correlations with CAP-2 (r=-0.849, p_uncorr_<0.001, p_BH_<0.001) and CAP-4 (r=-0.851, p_uncorr_<0.001, p_BH_<0.001). CAP-2 was found to be strongly anti-correlated with CAP-7 (r=-0.792, p_uncorr_<0.001, p_BH_<0.001). CAP-3 had a significant negative correlation with CAP-5 (r=-0.997, p_uncorr_<0.001, p_BH_<0.001), while CAP-4 was anti-correlated with CAP-6 (r=-0.751, p_uncorr_<0.001, p_BH_<0.001). Additionally, CAP-6 and CAP-7 exhibited a notable anti-correlation with each other (r=-0.786, p_uncorr_<0.001, p_BH_<0.001).

### Main Effect Time (Pre- vs. Post-Instruction)

A contrast analysis was performed on each linear mixed effect model to evaluate mean differences for each CAP metric from pre-instruction to post-instruction for each task (**Fig. 3**). These analyses were conducted by probing, for the main effect of time, several linear mixed-effects models (one model per temporal dynamic per CAP) that accounted for random effects due to individual differences while controlling for demographic variables including age, sex, ethnicity, household income, GPA, and years enrolled. For analyses involving CAP counts, the total number of TRs per participant was included as a covariate to account for differences in available data points due to motion scrubbing and task design. All p-values were corrected for multiple comparisons using the Benjamini-Hochberg procedure.

**Figure 3.**
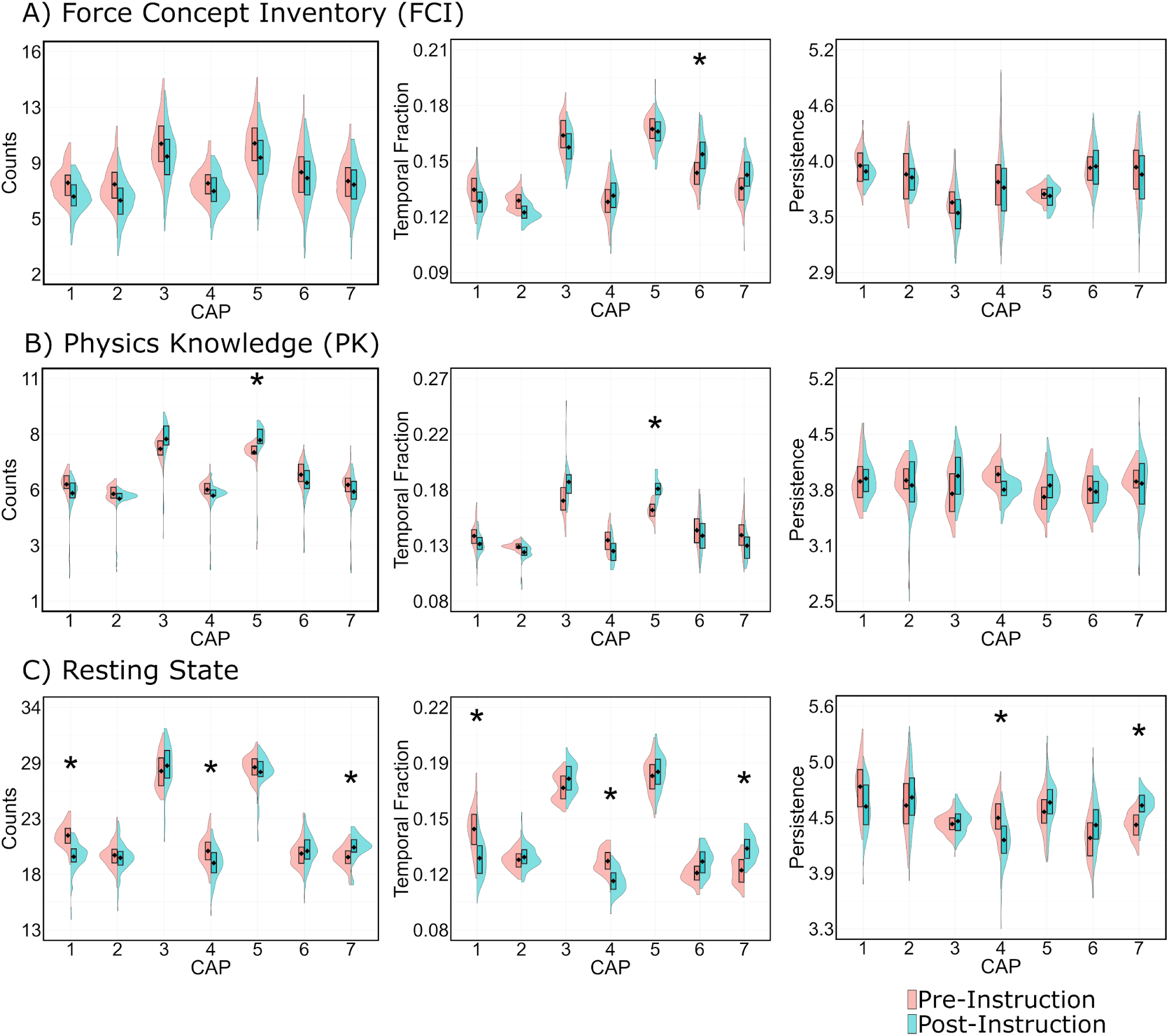
Main Effect of Time. Results of the main effect of time, which were probed from the linear mixed modeling analyses, are displayed for: A) Force Concept Inventory (FCI), B) Physics Knowledge (PK), and C) Resting State. Three CAP metrics are presented: Counts (left column), Temporal Fraction (middle column), and Persistence (right column). Each plot shows the distribution for each session (Pre-Instruction shown in pink and Post-Instruction in blue), accompanied by box plots displaying their means (black diamond) and edges representing the 25th and 75th quantile of the distribution. Asterisks (*) denote statistical significance of mean differences prior to Benjamini-Hochberg correction.

In the FCI task, only CAP-6 showed increased temporal fraction (Δ = 0.0140, *p*_*uncorr*_ = 0.018, *p*_*BH*_ = 0.191) from pre- to post-instruction. The PK task exhibited pre- to post-instruction changes with decreased temporal fraction (Δ = 0.019, *p*_*uncorr*_< 0.001, *p*_*BH*_= 0.067) and counts (Δ = 0.621, *p*_*uncorr*_= 0.031, *p*_*BH*_= 0.220) in CAP-5. The resting state showed the most widespread pre- to post-instruction changes, with significant alterations in three CAPs: CAP-1 showed decreased counts (Δ = −1.942, *p*_*uncorr*_< 0.001, *p*_*BH*_ < 0.001), temporal fraction (Δ = −0.019, *p*_*uncorr*_< 0.001, *p*_*BH*_ < 0.001), and persistence (Δ = −0.202, *p*_*uncorr*_= 0.036, *p*_*BH*_= 0.099), CAP-4 showed decreased counts (Δ = −1.090, *p*_*uncorr*_= 0.011, *p*_*BH*_= 0.039), temporal fraction (Δ = −0.012, *p*_*uncorr*_< 0.001, *p*_*BH*_ = 0.001), and persistence (Δ = −0.210, *p*_*uncorr*_= 0.025, *p*_*BH*_= 0.099), and CAP-7 showed increased counts (Δ = 1.130, *p*_*uncorr*_= 0.019, *p*_*BH*_= 0.044), temporal fraction (Δ = 0.014, *p*_*uncorr*_< 0.001, *p*_*BH*_ = 0.001), and persistence (Δ = 0.203, *p*_*uncorr*_= 0.047, *p*_*BH*_= 0.099). The delta change values and associated p-values for all tested models can be found in **Supplementary Table 4**.

### Main Effect of Instruction (Active Learning vs. Lecture-Based Classrooms)

Probing the linear mixed-models for the main effect of instruction revealed several significant effects. For the main effect of instruction **(Fig. 4)**, the active learning students exhibited lower persistence (Δ = − 0. 267, *p*_*uncorr*_= 0. 046, *p*_*BH*_ = 0. 322) of CAP-2 compared to the lecture-based students during the FCI task. Additionally, the active learning students also showed reduced persistence in CAP-2 (Δ = 0. 346, *p*_*uncorr*_ = 0. 014, *p*_*BH*_= 0. 047) and CAP-3 (Δ = 0. 394, *p*_*uncorr*_= 0. 007, *p*_*BH*_= 0.047) during the PK task when compared to the lecture-based students. The delta change values and associated p-values for all tested models can be found in **Supplementary Table 5**.

**Figure 4.**
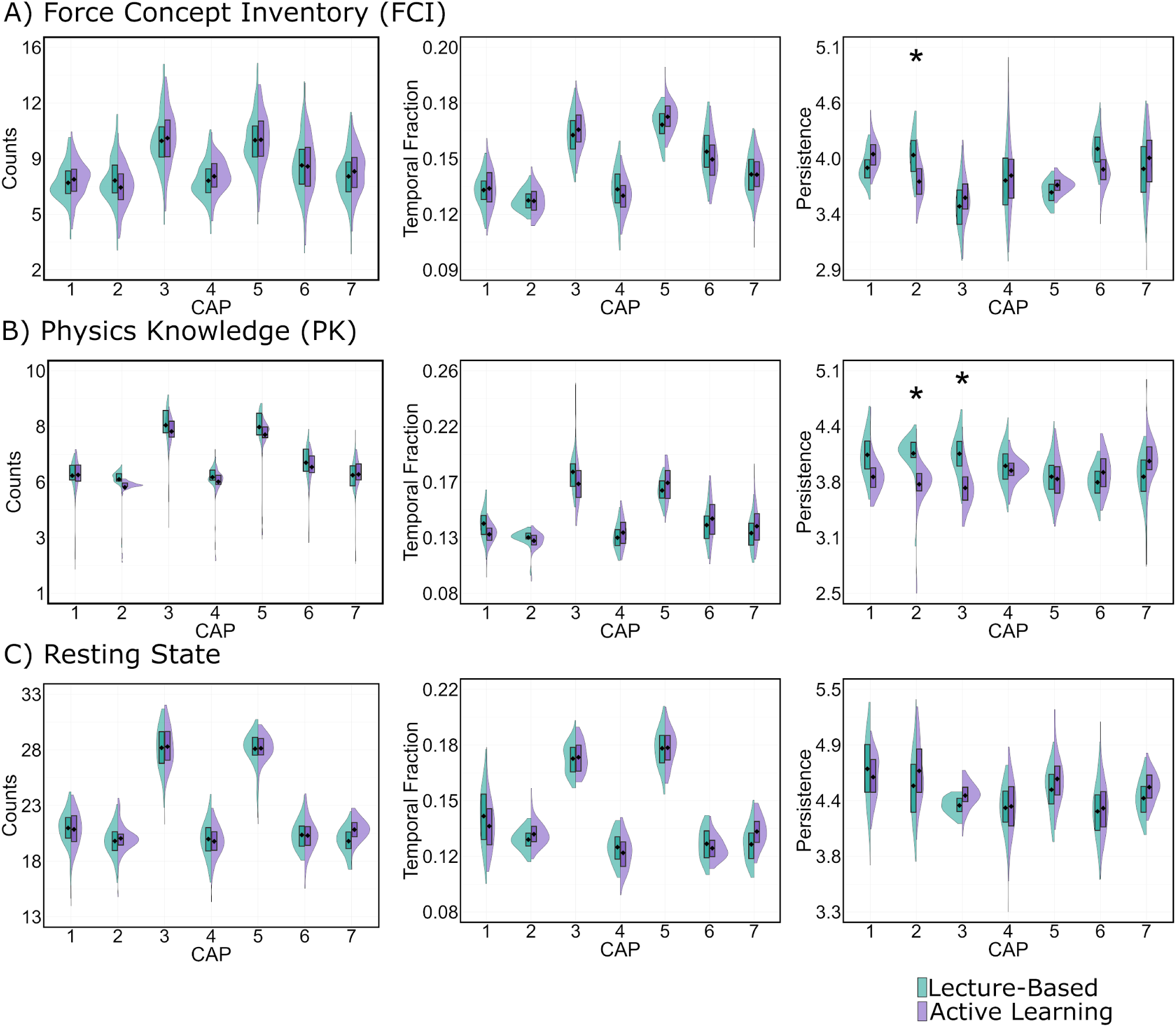
Main Effect of Instruction. Results of the main effect of instruction, which were probed from the linear mixed modeling analyses, are displayed for: A) Force Concept Inventory (FCI), B) Physics Knowledge (PK), and C) Resting State. Three CAP metrics are presented: Counts (left column), Temporal Fraction (middle column), and Persistence (right column). Each plot shows the distribution for each classroom (Active Learning shown in purple and Lecture-Based in green), accompanied by box plots displaying their means (black diamond) and edges representing the 25th and 75th quantile of the distribution. Asterisks (*) denote statistical significance of mean differences prior to Benjamini-Hochberg correction.

### Interaction Between Time and Instruction

Several significant interaction effects emerged (**Fig. 5**) from our linear mixed-effects models examining how instructional methods moderated changes in CAP metrics over time. These models included time (pre vs. post) and instruction type (active vs. lecture-based) as fixed effects, along with their interaction term, while controlling for demographic variables and random effects. The resulting beta coefficients represent how the change in CAP metrics from pre- to post-instruction differed between instructional groups, with positive values indicating greater increases or smaller decreases in temporal dynamics of a CAP within the active learning group compared to the lecture-based group.

**Figure 5.**
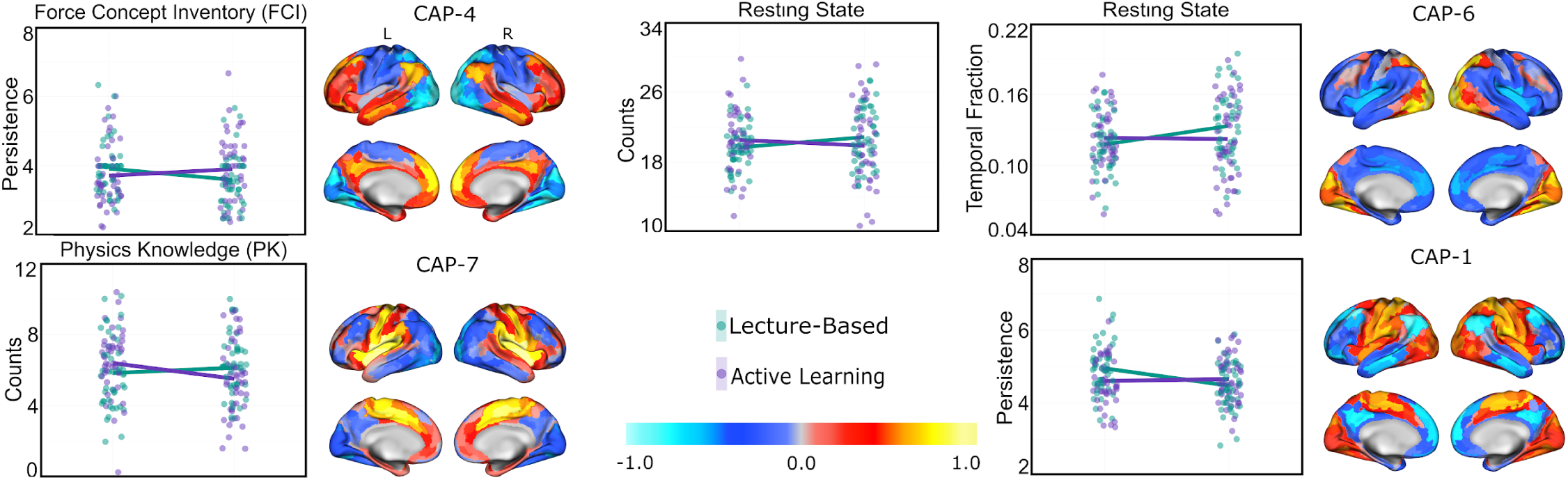
Interaction Effects. Interaction plots depicting changes in mean CAP counts, temporal fraction, and persistence from pre- to post-instruction for lecture-based (green) and active learning (purple) instructional methods across FCI, PK, and resting state data. Each plot represents a specific CAP (depicted in the right column) that showed a significant interaction effect prior to Benjamini-Hochberg correction: persistence of CAP-4 for FCI, counts of CAP-7 for PK, counts and temporal fraction of CAP-6, as well as counts of CAP-1, for resting state.

For the FCI task, CAP-4 showed a significant interaction with the active learning students demonstrating increased persistence (β = 0. 549, *p*_*uncorr*_= 0. 013, *p*_*BH*_= 0. 093) from pre- to post-instruction relative to the lecture-based student. For the PK task, the lecture-based students showed increased counts (β = −1. 120, *p*_*uncorr*_= 0. 031, *p*_*BH*_= 0. 215) from pre- to post-instruction compared to the active learning students in CAP-7. For resting state, from pre- to post-instruction, the active learning students demonstrated increased persistence (β = 0. 494, *p*_*uncorr*_= 0. 009, *p*_*BH*_= 0. 067) relative to the lecture-based instruction students in CAP-1 while the lecture-based instruction group exhibited increased counts (β = − 1. 860, *p*_*uncorr*_= 0. 0484, *p*_*BH*_= 0.338) and temporal fraction (β = − 0. 016, *p*_*uncorr*_ = 0. 037, *p*_*BH*_= 0.262) in CAP-6 when compared to the active learning students. The beta coefficients and associated p-values for all tested models can be found in Supplementary Table 6.

## Discussion

In this study, we investigated changes in the temporal dynamics of seven distinct CAPs, examining how different physics instructional methodologies modulate brain state dynamics over time. Drawing from embodied cognition theory and principles of Hebbian learning, we hypothesized that active learning and traditional lecture-based instructional approaches would differentially influence how students process and internalize physics concepts. Using three CAP metrics (i.e., counts, temporal fraction, and persistence), we evaluated changes in undergraduate students’ neural dynamics before and after a semester of introductory physics during both task-based (i.e., Force Concept Inventory (FCI) and physics knowledge (PK)) and resting state conditions. Our analysis revealed that the timing and type of instructional method and their interaction selectively modulate the temporal dynamics of distinct brain states. Supporting our hypotheses, these findings suggest that different instructional methodologies, specifically active and passive learning, result in divergent neural reconfigurations among students, potentially through differential engagement of sensorimotor and observational learning mechanisms.

A notable observation was that physics instruction led to significant changes in neural engagement, especially in the way students construct and utilize conceptual representations during problem solving. During the FCI task, we observed a notable increase in the temporal fraction of CAP-6, a pattern primarily characterized by activations in the Secondary Visual Network and the Dorsal Attention Network, at post-instruction. Given that the FCI task requires familiarity with Newtonian principles governing objects in motion and at rest, effective conceptual reasoning relies on constructing internal schemas^48^. These schemas allow students to form representations of potential solutions and the constraints of a problem, enabling them to quickly recognize patterns and draw upon prior knowledge to solve novel problems. During physics instruction, students encounter numerous problems related to Newtonian motion. Through repeated exposure, they develop strategies for identifying key visual features and constraints that determine an object’s trajectory. Furthermore, in contrast to novices, experts tend to spend more time constructing robust representations during the problem-solving process^49^. Consequently, at post-instruction, the increased engagement of the CAP associated with activations in the Secondary Visual Network, important for object recognition and mental simulation^50^, and the Dorsal Attention Network, critical for focusing attention on relevant features^51^, likely reflects students’ ability to generate more structured representations of motion problems and demonstrates expert-like thinking. This enhanced strategy supports the identification of features essential to applying Newtonian principles, thereby facilitating more effective problem-solving. Additionally, during the PK task, we observed an increase in both counts and temporal fraction within CAP-5 at post-instruction. These findings demonstrate the relevance of embodied cognition for physics-related semantic retrieval, as indicated by the involvement of the somatomotor network. In addition, these results are corroborated by prior findings from ^52^, which indicated that various physics concepts, such as those related to the causality of forces, flow of energy, and periodicity, were associated with activation of regions in the Somatomotor Network. Additionally, this CAP showed a more refined pattern of activation that also included the MT+ Complex and auditory association cortex. This CAP may be another reflection of increasing expertise as students develop more specialized neural representations that integrate visual motion processing and auditory associations, potentially drawing on visual recall of familiar diagrams or equations and auditory recall of verbal explanations^53,54^, while maintaining embodied components of physics knowledge.

In addition to task-related changes, our analyses revealed that the intrinsic organization of resting-state networks was reconfigured following physics instruction, suggesting a broader pattern of neural re-configuration^55^. Evidence suggests that learning interventions can induce changes in these resting-state patterns^56^. In line with our hypotheses, we observed significant changes from pre- to post-instruction in two complementary brain states: increased counts, temporal fraction, and persistence of CAP-7, characterized by Somatomotor and Cingulo-Opercular Network activation. This result suggests that physics learning, regardless of instructional methodology, heavily relies on embodied cognitive processes. Mason & Just^57^ demonstrated this relationship in their investigation of how naive participants learn physics concepts. In that study, participants viewed and learned about mechanical systems, such as a bathroom scale with its levers, springs, and mechanisms. Throughout the learning process, from initial encoding to forming mental representations and understanding system functionality, participants consistently engaged in somatomotor regions, highlighting the embodied nature of physics learning. Furthermore, the involvement of the Cingulo-Opercular network may be demonstrative of the importance of sustaining attention, while simultaneously monitoring for naive or incorrect beliefs^58^ about foundational physics concepts. Ultimately, the changes we observed in resting state dynamics in the present study align with principles of Hebbian learning and metastability. The repeated activation of these CAPs during physics learning appears to result in internal neural restructuring, potentially optimizing the brain for future acquisition of advanced physics concepts that require sensorimotor and visuospatial processing for initial comprehension^59^. This pattern suggests that initial physics concept learning relies heavily on sensorimotor processing, and the increased engagement of this CAP during post-instruction resting state may reflect neural restructuring that facilitates sensorimotor integration in physics learning. Interestingly, in contrast to our hypotheses, we observed decreased counts, temporal fraction, and persistence of CAP-4 from pre- to post-instruction. This CAP, characterized by increased activation in the Default Mode Network, Language Network, and Frontoparietal Network, most likely represents more effortful processing of physics concepts. However, this decline aligns with theories of skill acquisition and automaticity development^60,61^, which posit that initial learning stages require more controlled processing before transitioning to automatic processing, typically marked by reduced activation in control network regions as expertise develops. In the context of resting state activity, this reduction in CAP-4 engagement could reflect intrinsic cortical reconfiguration that primes the brain for more efficient physics concept processing^62^. Similarly, we observed decreased counts and temporal fraction of CAP-1 from pre- to post-instruction, a multimodal pattern showing significant correspondence with the Somatomotor Network, Visual Network, Cingulo-Opercular Network, and Dorsal Attention Network. While this CAP showed increased persistence in the active learning group relative to the passive learning group, this finding should be interpreted cautiously as the overall temporal dynamics of CAP-1 during resting state showed decreases across all temporal metrics, suggesting this interaction effect may be influencing the general pattern. One possible explanation may be due to the strong anticorrelation between CAP-1 and CAP-4, potentially indicating these patterns may operate in tandem, with decreases in effortful processing (i.e., CAP-4) leading to subsequent decreases in CAP-1 engagement at post-instruction.

Furthermore, beyond the general post-instruction learning effects, our findings revealed that different physics instruction methods significantly influenced the temporal dynamics of physics-related brain states, with distinct patterns emerging between lecture-based and active learning methodologies. During the FCI task, students in lecture-based environments exhibited decreased CAP-4 persistence from pre- to post-instruction, whereas those in active learning environments demonstrated enhanced persistence of this pattern. A critical requirement of the FCI task is the suppression of intuitive but non-Newtonian conceptualizations of motion trajectories^48^. CAP-4 shows its strongest positive activation in the anterior cingulate cortex (ACC), a region essential for cognitive control and particularly for suppressing competing conceptual frameworks^63^. This aligns with findings from Allaire-Duquette et al. ^64^ which demonstrate that physics experts consistently engage neural mechanisms to suppress intuitive physics concepts when contextually required. This suggests that active learning environments may facilitate the development of more expert-like neural patterns. Moreover, the ACC not only suppresses errors but also integrates past feedback to guide future decision-making processes^65^. Engagement of the posterior cingulate cortex and lateral temporal regions in CAP-4 provides additional insight into the learning process. These regions may support episodic memory integration, specifically the incorporation of experiential knowledge into conceptual understanding^66,67^. This activation pattern is particularly significant from the perspective of the embodied cognition framework. Active learning environments typically incorporate direct physical engagement with physics concepts through model manipulation and kinesthetic learning experiences. Consequently, the increased CAP-4 persistence, which entails sustained periods of integrating physical classroom experiences while simultaneously suppressing non-Newtonian intuitions, may reflect the effects of embodied learning aspects often employed in active learning contexts. Additionally, concurrent activation of the Dorsolateral Prefrontal Cortex and Orbitofrontal Cortex further supports this interpretation, as these regions are instrumental in integrating multiple information streams during decision-making processes^68–70^. This evidence suggests that the enhanced CAP-4 persistence observed in active learning students during FCI tasks reflects a sophisticated cognitive strategy that integrates embodied experience and active suppression of non-Newtonian intuitions during periods of self-referential thinking. Furthermore, prominent activation of the ACC and the auditory association areas positions the active learning group to engage in greater insight problem solving^71,72^, allowing students to potentially recognize unproductive cognitive strategies and transition into more effective ones. Consequently, such a brain state may suggest that active learning students approach physics-related conceptual reasoning in a fundamentally different manner compared to their lecture-based instruction counterparts. Evidence for this distinction emerged from the main effect of class in an FCI task, where the lecture-based group exhibited greater persistence of CAP-2, a pattern characterized by activations in the Default Mode and Frontoparietal networks. Although CAP-2 shares many overlapping active regions with CAP-4, it shows notably less anterior cingulate cortex (ACC) activation but substantially more activation in the inferior parietal cortex, a brain area significantly involved in visuospatial processing^73^. This pattern suggests that, when solving physics conceptual reasoning problems related to Newtonian motion, active learning students may rely more on their direct physical experiences with the concepts. By contrast, students taught primarily through lectures may rely more heavily on visuospatial processing strategies. Overall, these findings imply that students’ learning environments can meaningfully influence how they engage in and approach physics conceptual reasoning tasks.

The instructional methodology differences were also evident in the PK task, where the lecture-based group exhibited increased counts of CAP-7, characterized by primary activations in both the Cingulo-Opercular Network and Somatomotor Network, from pre- to post-instruction when compared to the active learning group. This finding aligns with previous research demonstrating that embodied cognition plays a fundamental role in physics learning, regardless of the instructional methodology used to present the information. Moreover, despite the involvement of embodied cognition, the increased counts of CAP-7 during physics-related semantic retrieval may also reflect the influence of lecture-based learning. Notably, CAP-7 exhibits prominent activation in the Insula and Frontal Operculum, implicated in language processing and multisensory integration^74,75^, and in the Early Auditory region, which is integral for processing speech^76^. Additionally, previous studies have demonstrated that the Cingulo-Opercular Network is involved in maintaining cognitive control for both word and speech recognition^58,77^. Given that lecture-based environments are typically instructor-driven and emphasize rote, recognition-based memorization^78^, CAP-7 suggests that while embodied cognition may be agnostic to instructional methodology, the chosen instructional approach may induce additional neurobiological changes that mirror the manner in which the content was disseminated. Specifically, lecture-based courses may require enhanced monitoring and sustained attention to external auditory cues, promoting the formation of internal, verbally mediated representations of physics concepts. During semantic retrieval, this may necessitate internal monitoring to ensure that the recognition of physics terms and their associated meanings remains consistent with prior lecture exposure and the associated sensorimotor experiences used to understand these concepts during learning. Furthermore, the main effect of class showed that lecture-based instruction groups displayed increased persistence of CAP-2 and CAP-3 compared to the active learning group. CAP-2 is characterized by activations in the Default Mode Network and Frontoparietal Network, while CAP-3 involves the Default Mode Network, Frontoparietal Network, and Cingulo-Opercular Network. Considering the high anticorrelation between CAP-2 and CAP-7, along with prominent activations in working-memory regions (Dorsolateral Prefrontal Cortex)^69^, autobiographical memory regions (Posterior Cingulate and Lateral Prefrontal Cortex)^66,67^, and visual word recognition (Inferior Parietal Lobule)^79^, CAP-2 may work in tandem with CAP-7, demonstrating a toggling between internal retrieval of semantic-related memories (i.e., CAP-2) and extended periods of external focus to correctly recognize specific physics terminology (i.e., CAP-7)^77^. Lastly, CAP-3 features activation in regions associated with working memory (Dorsolateral Prefrontal Cortex, Superior Parietal Lobule)^80^, visual processing (Superior Parietal Lobule, Inferior Parietal Lobule)^81^, language processing (Inferior Frontal)^82^, and recognition memory (Subcortical)^83,84^, which may further highlight how content is processed or encoded in lecture-based students. Considering that lecture-based instruction often emphasizes memorization and observational learning, these findings suggest a reliance on a more recognition and externally-based approach to physics-related semantic retrieval.

Finally, these instructional methodology differences were most prominently reflected in our resting state analyses, where, in direct support of our hypotheses, the active learning group exhibited a marked increase in the persistence of CAP-6 during resting state, from pre- to post-instruction, compared to the lecture-based instruction group. CAP-6 is a multimodal brain state characterized by activations across several networks, including the visual, dorsal attention, Cingulo-Opercular, and Somatomotor Networks. We posited that active learning would preferentially engage CAPs associated with sensorimotor processing, given that active learning STEM courses often employ more kinetic, hands-on approaches which, according to embodied cognition literature, should increase activation in sensorimotor regions^22,57^. Additionally, also aligned with our hypotheses, the lecture-based instruction group demonstrated increased counts and temporal fraction of CAP-6 from pre- to post-instruction during the PK task, reflecting that observation is one of their primary modes of learning. This aligns with research showing that observational learning activates visual processing regions and the superior parietal lobule^41,85^. The predominance of visual instruction in lecture-based settings appears to strengthen these visual processing networks. Both of these findings suggest a functional reorganization of the brain that becomes evident even at rest. Specifically, in the active learning group, this reorganization enhances sensorimotor integration, whereas in the lecture-based group, it enhances visual activation. These findings align with the principle of Hebbian learning, where persistent activations of certain network activation patterns, in response to repeated engagement with specific tasks, lead to intrinsic cortical restructuring and stable, recurrent state^27–29^. For both classes, such reorganization could promote improved comprehension of the physics material in the modality through which it is primarily delivered. In particular, for the active learning class, this reorganization may facilitate more efficient processing of physics concepts^62^, which typically requires substantial sensorimotor engagement during the learning process^57^. Notably, although CAP-6 declined in both temporal fraction and counts from pre- to post-instruction among all students, its relative preservation among active learning students likely reflects the distinct demands of their learning environment. Active learning courses require students to engage with physics content through multiple modalities simultaneously, specifically by physically manipulating objects, visually observing demonstrations, and mentally processing complex concepts, resulting in the reinforcement of sensorimotor-conceptual associations. Over time, such recurrent exposure and integration encourage long-term network reorganization that remains primed for reactivation to support future learning, potentially enabling a more cohesive comprehension of physics content or facilitating more effective absorption of material in the primary instructional mode.

The current study has several limitations. First, our original sample size was reduced from 121 to 90 participants due to our stringent exclusion criteria, which required at least one functional run of both resting state and task-based fMRI data for both pre- and post-instruction sessions; participants were excluded for various reasons including non-return for post-instruction scans, technical issues, MRIQC flagging for high mean framewise displacement, ghost-to-signal ratio, low signal-to-noise ratio, entropy focus criterion, or missing functional runs from one or more tasks. Importantly, t-tests and chi-square tests revealed no demographic differences between the active learning and lecture-based groups, and the final sample sizes were similar, with 46 active learning and 44 lecture-based participants. Second, our analytic approach identified shared CAPs across resting state and task-based fMRI data, which facilitated comparisons across instructional methods but did not allow for the detection of potential group-specific CAPs that might have emerged if clustering had been performed separately. Future studies may consider separate clustering approaches for each instructional group or task to better explore unique neural activation patterns. Finally, the exclusion of volumes with framewise displacement exceeding 0.35 mm was necessary to prevent high-motion artifacts from confounding the clustering results and ensure CAP integrity^86^, although this may have impacted temporal continuity metrics such as persistence; runs with mean framewise displacement in the 99th percentile were excluded, and participants included in the analysis had less than 40% of resting-state frames and 20% of task data frames scrubbed, aligning with prior CAP studies that retained data integrity without interpolating missing volumes^87–89^.

Overall, in this study, we examined how university physics learning acquired via different instructional methodologies (i.e., active versus passive learning) is associated with varying temporal dynamics of brain states. Our findings indicate significant intrinsic neural restructuring over a semester of learning. In line with our hypotheses, we found that physics learning across all students relied robustly on frontoparietal, somatomotor, and visual processing networks, which collectively support spatial comprehension and mathematical thinking. Specifically among active learning students, persistence of a somatomotor network (i.e., CAP-1) increased at post-instruction, supporting an embodied cognition framework. In contrast, lecture-based students exhibited greater persistence of a visuospatial network (i.e., CAP-6), suggesting a reliance on visual observation strategies. Moreover, additional evidence emerged, suggesting that embodied cognition is also utilized to an extent by lecture-based students when engaged in physics-related semantic retrieval (CAP-7), suggesting that embodied cognition is foundational to physics learning; however, the overall brain pattern among students may be significantly influenced by the instructional methodology used to disseminate information. Taken together, these findings provide evidence that the kinetic, hands-on approaches employed in active learning classrooms yield measurable differences in brain activation compared to traditional lecture-based instruction. These results enhance our understanding of how university physics education influences brain dynamics. This work indicates that instructors’ pedagogical choices influence the neural strategies that students employ during physics cognition, ultimately playing a role in reshaping the neural architecture of learning.

## Methods

### Participants

The sample consisted of 121 healthy, right-handed, undergraduate students (mean age=19.8 ± 1.5, range=18-26 years; 55 females) enrolled in a 15-week semester of calculus-based, introductory physics course at Florida International University (FIU). Students received either traditional, lecture-based instruction (*n*=60) or Modeling Instruction (*n*=61), an active learning approach that engages students in model building and collaborative group activities to understand physics concepts^42^. At enrollment, participants provided demographic information, including age, sex, ethnicity, household income, grade point average (GPA), and years enrolled at FIU (i.e., freshman, sophomore, junior, or senior) (**Table 1**) (**Supplementary Table 1**). All participants were free from cognitive, neurological, or psychiatric conditions and reported no use of psychotropic medications.

**Table 1.** Participant Demographic Information.

|  | Modeling Instruction (n=61) |  | Lecture-Based Instruction (n=60) |  |
| --- | --- | --- | --- | --- |
|  | N | Percentage | N | Percentage |
| **Gender** |  |  |  |  |
| Male | 37 | 61 | 31 | 52 |
| Female | 24 | 39 | 29 | 48 |
| **Ethnicity** |  |  |  |  |
| Hispanic | 44 | 72 | 43 | 72 |
| Non-Hispanic | 17 | 28 | 17 | 28 |
| **Household Income** |  |  |  |  |
| < \$15,000 | 12 | 20 | 13 | 22 |
| \$15,000 - \$34,999 | 10 | 16 | 16 | 27 |
| \$35,000 - \$49,999 | 6 | 10 | 9 | 15 |
| \$50,000 - \$74,999 | 15 | 25 | 7 | 12 |
| \$75,000 - \$99,999 | 11 | 18 | 6 | 10 |
| > \$100,000 | 7 | 11 | 9 | 15 |
| **Years Enrolled** |  |  |  |  |
| Freshman | 6 | 10 | 6 | 10 |
| Sophomore | 27 | 44 | 29 | 48 |
| Junior | 21 | 34 | 18 | 30 |
| Senior | 7 | 11 | 7 | 12 |
|  | **Grand Mean** | **Std. Dev.** | **Grand Mean** | **Std. Dev.** |
| **Age** | 19.84 | 1.67 | 19.73 | 1.43 |
| **GPA** | 3.29 | 0.67 | 3.29 | 0.38 |

Note. "N" represents the sample size of the group, and "Percentage" represents the percentage of participants in that group for each categorical variable (i.e., gender, ethnicity, household income, and years enrolled).

### Procedures

Behavioral and neuroimaging data collection occurred during two time periods: pre- and post-instruction. Multiple different recruitment strategies were employed, including several in-class visits (at the professor’s discretion), emailing eligible students, and distributing study flyers across FIU’s campus. Enrolled participants completed an fMRI session at the start of the course (i.e., pre-instruction), no later than the fourth week of instruction and before the first exam. Neuroimaging sessions were conducted off-campus and participants were provided with free parking and/or FIU-organized transportation to and from the MRI site. At the end of the semester, participants completed their post-instruction data collection session no later than two weeks after the semester concluded and after final exams were completed. Written informed consent was obtained in accordance with FIU’s Institutional Review Board approval. Additionally, participants were monetarily compensated for their study participation.

### MRI Data Acquisition

MRI data were acquired on a GE 3T Healthcare Discovery 750W MRI scanner at the University of Miami. Functional imaging data were acquired with an interleaved gradient-echo, echo planar imaging (EPI) sequence (repetition time [TR]/echo time [TE] = 2000/30ms, flip angle = 75°, field of view (FOV) = 220x220mm, matrix size = 64x64, voxels dimensions = 3.4×3.4×3.4mm, 42 axial oblique slices). T1-weighted structural data were also acquired using a 3D fast spoiled gradient recall brain volume (FSPGR BRAVO) sequence with 186 contiguous sagittal slices (TI = 650ms, bandwidth = 25.0kHz, flip angle = 12°, FOV = 256x256mm, and slice thickness = 1.0mm).

### fMRI Tasks

#### Conceptual Reasoning Task

Participants completed a self-paced, conceptual reasoning task consisting of questions derived from the Force Concept Inventory (FCI) (**Fig. 6A-C**) ^90–92^ that was adapted for the MRI environment^48^. FCI trials included textual descriptions and illustrations of scenarios, including objects at rest or in motion, accompanied by one correct Newtonian solution and several incorrect non-Newtonian solutions. Control trials included similar visual stimuli to FCI trials but tested general reading comprehension and/or shape discrimination. Each trial was presented in blocks composed of three sequential view screens: Phase I: Scenario, which included an illustration and text describing a physical scenario; Phase II: Question, which involved presenting a question related to the scenario; and Phase II: Answer, where participants were presented with four answer choices. Both FCI and control blocks had a maximum duration of 45 sec and were followed by a fixation cross with a minimum duration of 10 sec. The total duration for each FCI run was 5 min 44 sec; data were collected during three runs for a total duration of ∼16 minutes, resulting in 170 volumes being collected per participant.

**Figure 6.**
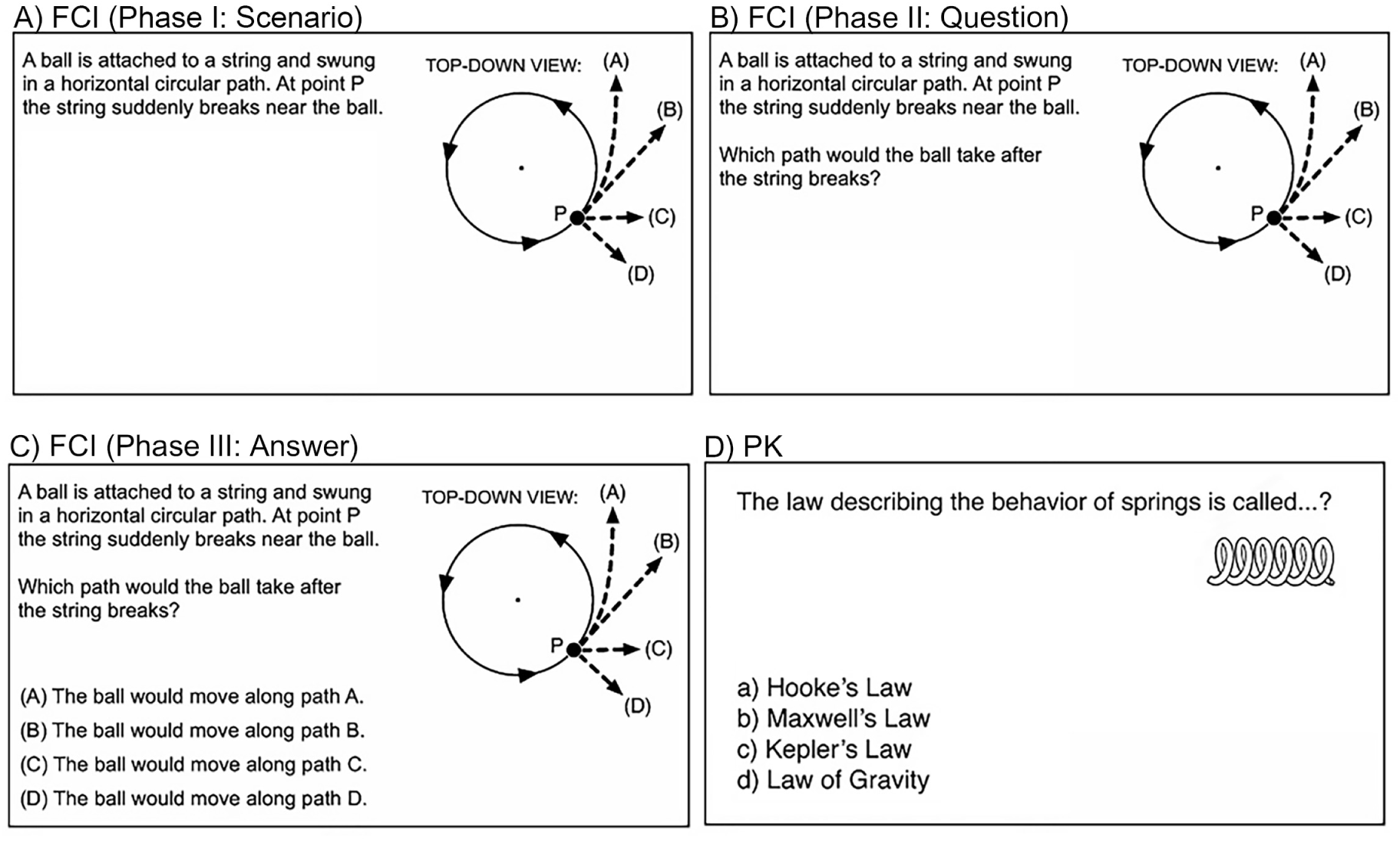
Force Concept Inventory (FCI) and Physics Knowledge (PK) Tasks. Example items of the in-scanner physics tasks, including the three phases of the FCI task (A) Phase I: Scenario, (B) Phase II: Question, (C) Phase III: Answer, and (D) the PK task.

#### Knowledge Retrieval Task

Participants also completed the physics knowledge (PK) task (**Fig. 6D**), which was adapted from a general knowledge task of semantic retrieval ^93^. The PK task was presented in a block design and probed for brain activation associated with physics-based content knowledge. Students viewed physics questions related to general physics knowledge and their corresponding answer choices. A control condition was presented in which students viewed general knowledge questions with corresponding answer choices. PK and control blocks were 28 seconds long and included four questions per block (6.5 sec per question followed by 0.5 sec of quick fixation). Three blocks of physics or general questions (six question blocks total) were alternated with 10 sec of fixation. The total duration of one run was 4 min 2 sec; data were collected during two runs for a total duration of ∼8 minutes, resulting in 180 volumes being collected per participant.

#### Resting State

Participants were instructed to lie quietly with their eyes closed and not to fall asleep. The total duration of the resting state session was 12 minutes, resulting in 365 volumes of data being collected per participant. The first five volumes for each participant were removed to ensure only steady-state data were used in subsequent analyses, leaving 360 volumes of resting state data per participant.

### MRI Preprocessing

All functional runs within this study (i.e., FCI, PK, rest) were preprocessed independently. Each participant’s T1-weighted images were corrected for intensity non-uniformity with `ANT’s N4BiasFieldCorrection` tool^94,95^. Anatomical and functional images underwent additional preprocessing steps using `fMRIPrep` (v.1.5.0rc1)^96^. The T1-weighted (T1w) reference, which was used throughout the pipeline, was generated after T1w images were corrected for intensity non-uniformity with `ANT’s N4BiasFieldCorrection`. `Freesurfer’s mri_robust_template` was used to generate a T1w reference, which was used throughout the entire pipeline^95,97^. `Nipype’s` implementation of `ANT’s antsBrainExtraction` workflow was used to skullstrip the T1w reference using `OASIS30ANTs` as the target template ^98^. `FSL’s FAST` was used for brain tissue segmentation of the cerebrospinal fluid (CSF), white matter (WM), and gray matter (GM); brain surfaces were reconstructed using `Freesurfer’s recon_all`^99,100^. Preprocessing of functional images began with selecting a reference volume and generating a skullstripped version using a custom methodology of `fMRIPrep`. `Freesurfer’s bbregister`, which uses boundary-based registration, was used to coregister the T1w reference to the BOLD reference. The BOLD time series were then resampled onto surfaces of fsaverage5 space and resampled onto their original, native space by applying a single, composite transform to correct for head motion and susceptibility distortions. Additionally, the BOLD time series were high pass filtered, using a discrete cosine filter with a cutoff of 128s^101^. For each volume, except the first, the framewise displacement (FD) was computed. Nuisance signals from CSF, WM, and whole brain masks were extracted by using a set of physiological regressors, which were extracted to allow for both temporal component-based noise correction (tCompCor) and anatomical component-based noise correction (aCompCor)^102^. Additionally, the confound time series derived from head motion estimates were expanded to include its temporal derivatives and quadratic terms, resulting in a total of 24 head motion parameters (i.e., six base motion parameters, six temporal derivatives of six motion parameters, 12 quadratic terms of six motion parameters, and their six temporal derivatives). Estimates for the global, cerebrospinal fluid, and white matter signals were expanded to include their temporal derivatives and quadratic terms, resulting in a total of 12 signal-based parameters (i.e., three base signal parameters, three temporal derivatives of the three base parameters, the three quadratic terms of the base parameters, and the three quadratic terms of the temporal derivatives). Finally, all 24 head motion confound estimates, three high pass filter estimates, and a variable number of aCompCor estimates (components that explain the top 50% of the variance) were outputted into a tsv file to be used for later denoising steps^103^. An expanded description of the preprocessing workflow can be found in the Supplemental Information.

### Quality Control Assessment

Prior to time series extraction, `MRIQC`^104^ was used to calculate quality metrics for each participant’s functional runs, including mean framewise displacement, ghost-to-signal ratio, signal-to-noise ratio, and entropy focus criterion (an indicator of ghosting and blurring resulting from head motion)^105^. Based on the `MRIQC` reports, participants’ functional runs with a mean framewise displacement, ghost-to-signal ratio, or entropy focus criterion above the 99th percentile, or a signal-to-noise ratio below the 1st percentile, were excluded from the analysis. This exclusion criterion resulted in the removal of 17 participants from the resting state data, two participants from the FCI task, and one participant from the PK task. It is important to note that these exclusions pertained to participants where all functional runs from both fMRI sessions (i.e., pre- and post-instruction) were flagged by `MRIQC`. Additionally, the participants excluded from the FCI and PK tasks overlapped with those excluded from the resting state data.

### Denoising and Parcellation

Following quality control assessment, all runs and sessions from both the resting state and task data underwent identical denoising strategies. This uniformity was essential to ensure that the subsequent clustering of time points from the combined dataset was not influenced by variations in denoising approaches. Specifically, the time series of each participant’s task and resting state data were bandpass filtered (0.008–0.1 Hz), detrended, and subjected to nuisance regression. This regression involved orthogonalizing the time series against the six head motion parameters and their first derivatives, the global signal and its first derivative, and the first six aCompCor components derived from the white-matter and cerebrospinal fluid masks. Spatial dimensionality reduction was performed using the Human Connectome Project extended (HCPex) atlas, which is both a volumetric version and extension of Glasser et al.^44^ that incorporates an additional 66 subcortical areas resulting in a total of 426 nodes ^43^. Additionally, Huang et al.^43^ classified these 426 nodes into 23 cortical and subcortical divisions (**Fig. 7**).

**Figure 7.**
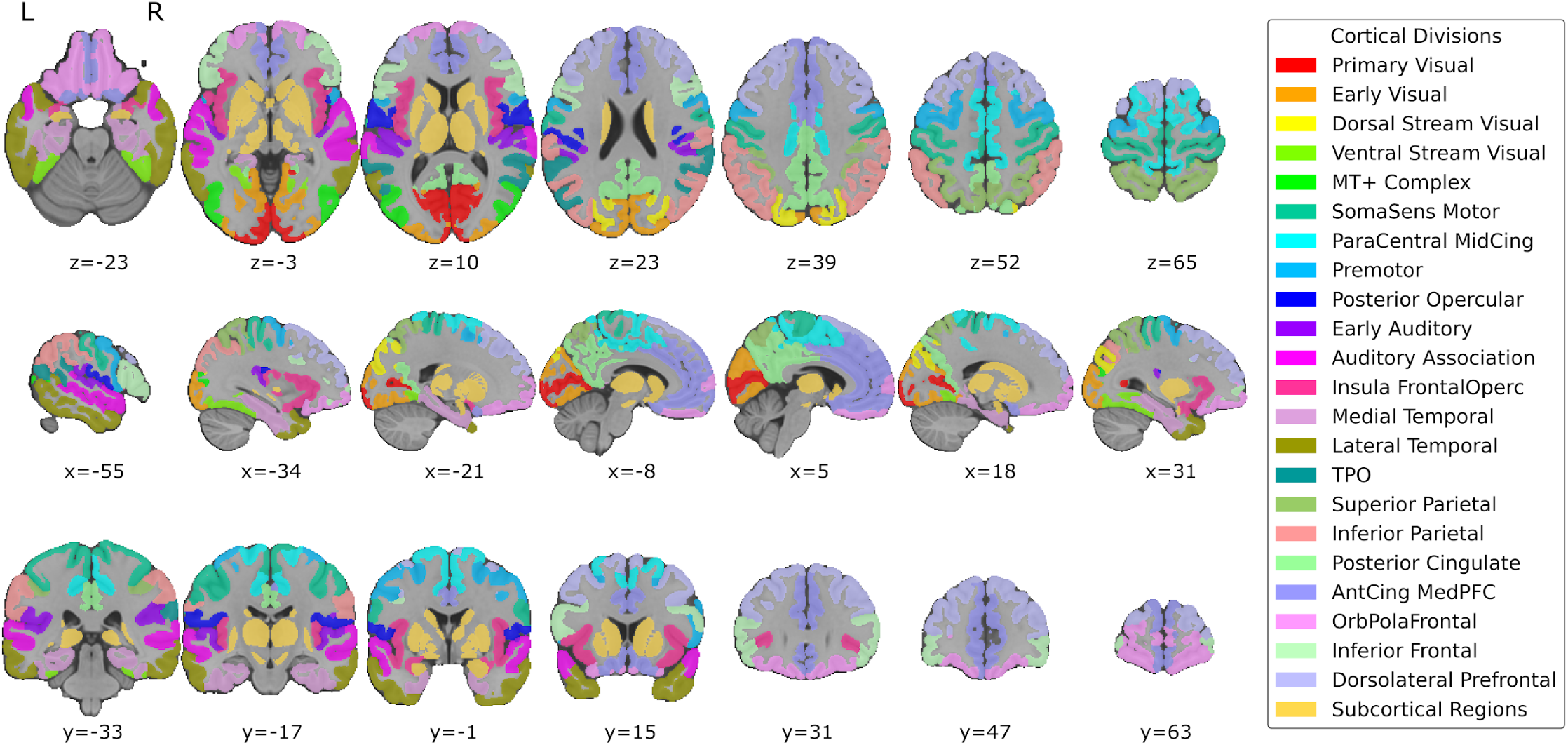
Cortical and Subcortical Divisions of the HCPex Atlas. The HCPex Atlas, which contains 426 nodes, was used to parcellate each participant’s preprocessed fMRI data. All 426 nodes within the atlas were classified into 23 cortical subdivisions, as described by Huang et al.^43^.

For task data, only volumes corresponding to the physics trials were retained (i.e., control trials were removed), resulting in approximately 40 and 45 frames retained per run for the FCI and PK tasks, respectively. To account for the hemodynamic delay^106^, onset times for each physics trial block were shifted forward by 4 seconds ^107^. Additionally, for both the physics task and resting state data, volumes from each functional run with a framewise displacement exceeding 0.35 mm were scrubbed. These TRs were removed to prevent clustering from being significantly affected by time points dominated by motion artifacts. Participants were also assessed to ensure that the proportion of scrubbed volumes did not exceed 20% per run for the physics task data and 40% per run for the resting state data. A stricter retention threshold was applied to the task data due to the difference in the number of frames being assessed (i.e., 40 and 45 frames per run for the FCI and PK tasks, respectively compared to 360 frames for resting state. These retention thresholds align with previous, recent CAP studies that retained participants with less than 30-50% of scrubbed data^87–89^. Afterwards, standardization was conducted within runs to allow the subsequent clustering procedure to prioritize patterns of network activation in all HCPex regions of interest (ROI) as opposed to the absolute magnitude differences, especially between task and rest data.

### Coactivation Pattern (CAP) Analysis

Once denoising was completed, coactivation pattern (CAP) analysis was performed. To this end, participants with at least one functional run across all sessions and tasks (*n*=90) were concatenated into a single matrix (rows: *N*_*participants*_ \**N*_*Repetition Time*_ ; columns: *N*_*ROIs*_). This criterion was applied to mitigate bias in clustering solutions and subsequent statistical analyses that could arise from the overrepresentation of data from specific sessions or tasks, or from participants of certain demographics contributing data exclusively to particular tasks and sessions. This approach allows for better identification of CAPs and statistical trends that are more broadly generalizable across all tasks and sessions. The matrix was then subjected to k-means clustering using Euclidean distance, where values of *k* ranging from 2 to 20 were tested. The optimal value of cluster size was determined using the elbow criterion, a heuristic where k is the value where the within-cluster sum of squares (i.e., inertia) stops decreasing exponentially^108^.

After the optimal value of *k* was determined, the cluster centroids, representing the average of all data points assigned to the same CAP, were extracted to generate five visualizations: a regional heatmap, a correlation matrix, surface plots for each CAP, and radar plots for each CAP. The regional heatmap was created by extracting each cluster centroid, calculating the average activation among all nodes assigned to the same cortical division within the HCPex atlas, as described by Huang et al. ^43^ and visualizing these averages. The correlation matrix was obtained by calculating the Pearson correlation between all possible pairs of CAPs, with p-values generated for each element in the matrix, to assess the linear relationship between each pair of CAPs. Surface plots were created by replacing each label in the HCPex atlas with its corresponding value from the cluster centroid and converting the atlas from Montreal Neuroimaging Institute (MNI) volumetric space into FreesurferLR (fsLR) surface space for visualization. Radar plots for each CAP were constructed by generating separate activation vectors for the “High Amplitude” activations, which are values above zero, and the “Low Amplitude” activations, which are values below zero. The cosine similarity between each activation vector and a binary representation of each *a priori* cortical division in the HCPex atlas^89,109,110^, was computed using the following formula:

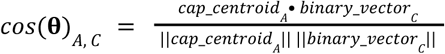

*w*ℎ*ere A* ∈ {*Hig*ℎ *Amplitude*, *Low Amplitude*} *and C* ∈ {*Cortical Division*_1_, *Cortical Division*_2_, …}

Each element within the CAP cluster centroid represented the relative activation or deactivation of an ROI belonging to one HCPex cortical division. Hence, the binary vectors representing the cortical divisions consisted of ‘1’s for the ROIs within the cluster centroid assigned to the same cortical division, while ‘0’s indicated otherwise. Furthermore, the absolute values of the negative activations within the cluster centroid were computed to restrict the “High Amplitude” and “Low Amplitude” cosine similarities from 0 to 1 for visualization purposes. Cosine similarities greater than 0.2 or less than −0.2 are reported in the results section.

Next, a network correspondence analysis was performed using the `cbig_network_correspondence` Python package ^46^. This analysis evaluated the overlap between positive activations of each CAP (with cosine similarity ≥ 0. 20) and the Cole-Antecevic atlas, which maps ROIs from the Glasser et al.^44^ atlas onto 12 distinct resting state networks ^45^. The package employs spin permutation methods^47^ to assess overlap significance, quantified through the dice coefficient. Finally, the CAP assignments of each participant’s time points for all runs and sessions of their task and resting state data were extracted to calculate three CAP measures for each session: **temporal fraction** (i.e., the proportion of time occupied by a single CAP), **persistence** (i.e., the average duration a CAP persists before transitioning to another CAP), and **counts** (i.e., the frequency of CAP initiation across the entire scan)^38^. Furthermore, to control the number of statistical comparisons for each task (specifically the FCI and PK tasks), multiple runs were treated as a single, continuous run for the calculation of each CAP metric ^111^.

### Statistical Analyses

We first assessed demographic differences between active learning and lecture-based instruction groups. Independent samples t-tests were used to compare age and GPA (both pre- and post-instruction), while chi-square tests were used to compare sex, ethnicity, household income, and years enrolled at FIU. These analyses were conducted for both the full sample (n=121) and the subsample of participants who had at least one functional run for all tasks and sessions (n=90). Next, using the calculated CAP metrics (i.e., temporal fraction, counts, and persistence), we assessed the main effects of time (i.e., pre- versus post-instruction), the main effects of instructional methodology (i.e., modeling versus lecture-based instruction), and the interaction effects between time and instructional methodology. Linear mixed-effects models were employed using the `lme4` package ^112^, regressing each CAP metric on the interaction between time and instructional methodology, while controlling for nuisance variables such as age, sex, ethnicity, household income, GPA, and number of years enrolled at FIU, and accounting for random effects due to individual differences. Furthermore, counts are raw frequency calculations of CAP initiations, unlike temporal fraction and persistence, which includes some level of accounting for differences in the magnitude of TRs due to being proportion and average duration-based metrics, respectively. Therefore, the total number of TRs per participant for the given task being analyzed was included as a covariate in all linear mixed-effects models assessing counts. This was done to ensure that observed differences in the counts of any CAP could be attributed to effects of interest as opposed to differences in the total number of TRs (as a consequence of scrubbing or the self-paced nature of the FCI task). Further descriptive information regarding the total number of TRs for contrasts of interest can be found in Supplemental Information. Finally, all linear mixed-effects models were probed for the main effects of time and instructional methodology using `R’s emmeans` ^113^ as the beta coefficients for time and class represent mean differences from the reference group, rather than capturing differences among the variable itself. To account for multiple comparisons, the Benjamini-Hochberg procedure (BH) was applied, and findings with corrected and uncorrected p-values are reported and discussed.

## Acknowledgments

Data collection for this project was funded by NSF REAL DRL-1420627 (ARL, EB, SMP, JEB). Contributions from co-authors were provided with support from NSF 1631325 (ARL, MCR, TS), NIH R01-DA041353 (ARL, MTS, MCR), NIH U01-DA041156 (ARL, MTS, MCR, KLB). Additional support was provided by the Office of Research and Economic Development and the Dissertation Fellowship from the University Graduate School at Florida International University (FIU). Special thanks to the FIU Instructional & Research Computing Center (IRCC, http://ircc.fiu.edu) for providing the HPC and computing resources that contributed to the research results reported within this paper, as well as to the Department of Psychology of the University of Miami for providing access to their MRI scanner. Additional thanks to Dr. Jeremy Elman for sharing semantic retrieval task stimuli. Lastly, the authors would like to thank the FIU undergraduate students who volunteered, participated, and contributed to this project.

## Data Availability

All tabular (behavioral) data used to perform the statistical analyses are available on the Open Science Framework (OSF) project page at https://osf.io/s6fp9/. Statistical brain volumes resulting from neuroimaging analyses (i.e., CAPs maps) are available at https://neurovault.org/collections/19622/.

## Code Availability

The source code for `neurocaps` ^114^, an open-source Python package developed for this study is available at https://github.com/donishadsmith/neurocaps. This package was used for time series extraction, clustering and identification of CAPs, and visualization of CAPs (i.e., heatmap, surface plots, correlation matrix, and cosine similarity plots). Additionally, the source code for the Network Correspondence Toolbox (NCT), which was used to identify the networks of the positive activations of each CAP, is available at https://github.com/rubykong/cbig_network_correspondence. Furthermore, the HCPex parcellation, scripts from the statistical analyses, and figures used in this study are available on the OSF project page at https://osf.io/s6fp9/.

## Author Contributions

DDS and ARL conceived and designed the study. JEB acquired data. DDS analyzed the data. DDS, JAP, KLB, MCR, and TS contributed scripts, pipelines, and figures. JSN and LQU provided analytic guidance and support for the CAP analysis. DDS and ARL wrote the paper, and all authors contributed to the revisions and approved the final version.

## References

1. Bao, L. & Koenig, K. Physics education research for 21st century learning. Discip. Interdiscip. Sci. Educ. Res. 1, 2 (2019).

2. Hoskinson, A.-M., Caballero, M. D. & Knight, J. K. How Can We Improve Problem Solving in Undergraduate Biology? Applying Lessons from 30 Years of Physics Education Research. CBE—Life Sci. Educ. 12, 153–161 (2013).

3. Dancy, M. et al. Physics instructors’ knowledge and use of active learning has increased over the last decade but most still lecture too much. Phys. Rev. Phys. Educ. Res. 20, 010119 (2024).

4. Freeman, S. et al. Active learning increases student performance in science, engineering, and mathematics. Proc. Natl. Acad. Sci. 111, 8410–8415 (2014).

5. Ješková, Z. et al. Active Learning in STEM Education with Regard to the Development of Inquiry Skills. Educ. Sci. 12, 686 (2022).

6. Minhas, P. S., Ghosh, A. & Swanzy, L. The effects of passive and active learning on student preference and performance in an undergraduate basic science course. Anat. Sci. Educ. 5, 200–207 (2012).

7. Chin, D. B., Chi, M. & Schwartz, D. L. A comparison of two methods of active learning in physics: inventing a general solution versus compare and contrast. Instr. Sci. 44, 177–195 (2016).

8. Commeford, K., Brewe, E. & Traxler, A. Characterizing active learning environments in physics using network analysis and classroom observations. Phys. Rev. Phys. Educ. Res. 17, 020136 (2021).

9. Dou, R., Brewe, E., Potvin, G., Zwolak, J. P. & Hazari, Z. Understanding the development of interest and self-efficacy in active-learning undergraduate physics courses. Int. J. Sci. Educ. 40, 1587–1605 (2018).

10. Durk, J., Davies, A., Hughes, R. & Jardine-Wright, L. Impact of an active learning physics workshop on secondary school students’ self-efficacy and ability. Phys. Rev. Phys. Educ. Res. 16, 020126 (2020).

11. Prada Nuñez, R., Hernández, C. A. & Gamboa, A. A. Active learning and knowledge in physics: a reading from classroom work. J. Phys. Conf. Ser. 1981, 012007 (2021).

12. Chi, M. T. H. & Wylie, R. The ICAP Framework: Linking Cognitive Engagement to Active Learning Outcomes. Educ. Psychol. 49, 219–243 (2014).

13. Georgiou, H. & Sharma, M. D. Does using active learning in thermodynamics lectures improve students’ conceptual understanding and learning experiences? Eur. J. Phys. 36, 015020 (2015).

14. Yannier, N., Hudson, S. E. & Koedinger, K. R. Active Learning is About More Than Hands-On: A Mixed-Reality AI System to Support STEM Education. *Int*. J. Artif. Intell. Educ. 30, 74–96 (2020).

15. Anderson, M. L. Embodied Cognition: A field guide. Artif. Intell. 149, 91–130 (2003).

16. Collins, J. A. & Olson, I. R. Knowledge is power: How conceptual knowledge transforms visual cognition. Psychon. Bull. Rev. 21, 843–860 (2014).

17. Borghi, A. M. & Cimatti, F. Embodied cognition and beyond: Acting and sensing the body. Neuropsychologia 48, 763–773 (2010).

18. Boulenger, V., Hauk, O. & Pulvermuller, F. Grasping Ideas with the Motor System: Semantic Somatotopy in Idiom Comprehension. Cereb. Cortex 19, 1905–1914 (2009).

19. Kontra, C., Lyons, D. J., Fischer, S. M. & Beilock, S. L. Physical Experience Enhances Science Learning. Psychol. Sci. 26, 737–749 (2015).

20. Moseley, R. L. & Pulvermüller, F. What can autism teach us about the role of sensorimotor systems in higher cognition? New clues from studies on language, action semantics, and abstract emotional concept processing. Cortex 100, 149–190 (2018).

21. Ptak, R., Doganci, N. & Bourgeois, A. From Action to Cognition: Neural Reuse, Network Theory and the Emergence of Higher Cognitive Functions. Brain Sci. 11, 1652 (2021).

22. Doganci, N., Iannotti, G. R., Coll, S. Y. & Ptak, R. How embodied is cognition? fMRI and behavioral evidence for common neural resources underlying motor planning and mental rotation of bodily stimuli. Cereb. Cortex 33, 11146–11156 (2023).

23. Kao, C.-H. et al. Functional brain network reconfiguration during learning in a dynamic environment. Nat. Commun. 11, 1682 (2020).

24. Lurie, D. J. et al. Questions and controversies in the study of time-varying functional connectivity in resting fMRI. Netw. Neurosci. 4, 30–69 (2020).

25. Hutchison, R. M. et al. Dynamic functional connectivity: Promise, issues, and interpretations. NeuroImage 80, 360–378 (2013).

26. Jiang, F. et al. Time-varying dynamic network model for dynamic resting state functional connectivity in fMRI and MEG imaging. NeuroImage 254, 119131 (2022).

27. Bassett, D. S. et al. Dynamic reconfiguration of human brain networks during learning. Proc. Natl. Acad. Sci. 108, 7641–7646 (2011).

28. Dosenbach, N. U. F. et al. Distinct brain networks for adaptive and stable task control in humans. Proc. Natl. Acad. Sci. 104, 11073–11078 (2007).

29. Harmelech, T., Preminger, S., Wertman, E. & Malach, R. The Day-After Effect: Long Term, Hebbian-Like Restructuring of Resting-State fMRI Patterns Induced by a Single Epoch of Cortical Activation. J. Neurosci. 33, 9488–9497 (2013).

30. Alderson, T. H., Bokde, A. L. W., Kelso, J. A. S., Maguire, L. & Coyle, D. Metastable neural dynamics underlies cognitive performance across multiple behavioural paradigms. Hum. Brain Mapp. 41, 3212–3234 (2020).

31. Deco, G., Kringelbach, M. L., Jirsa, V. K. & Ritter, P. The dynamics of resting fluctuations in the brain: metastability and its dynamical cortical core. Sci. Rep. 7, 3095 (2017).

32. Denkova, E., Nomi, J. S., Uddin, L. Q. & Jha, A. P. Dynamic brain network configurations during rest and an attention task with frequent occurrence of mind wandering. Hum. Brain Mapp. 40, 4564–4576 (2019).

33. Liu, X., Zhang, N., Chang, C. & Duyn, J. H. Co-activation patterns in resting-state fMRI signals. NeuroImage 180, 485–494 (2018).

34. Rabany, L. et al. Dynamic functional connectivity in schizophrenia and autism spectrum disorder: Convergence, divergence and classification. NeuroImage Clin. 24, 101966 (2019).

35. Cao, H. et al. Cross-paradigm connectivity: reliability, stability, and utility. Brain Imaging Behav. 15, 614–629 (2021).

36. Elliott, M. L. et al. General functional connectivity: Shared features of resting-state and task fMRI drive reliable and heritable individual differences in functional brain networks. NeuroImage 189, 516–532 (2019).

37. McCormick, E. M., Arnemann, K. L., Ito, T., Hanson, S. J. & Cole, M. W. Latent functional connectivity underlying multiple brain states. Netw. Neurosci. 6, 570–590 (2022).

38. Yang, H. et al. Reproducible coactivation patterns of functional brain networks reveal the aberrant dynamic state transition in schizophrenia. NeuroImage 237, 118193 (2021).

39. Bartley, J. E. et al. Meta-analytic evidence for a core problem solving network across multiple representational domains. Neurosci. Biobehav. Rev. 92, 318–337 (2018).

40. Cetron, J. S. et al. Using the force: STEM knowledge and experience construct shared neural representations of engineering concepts. Npj Sci. Learn. 5, 6 (2020).

41. Bellebaum, C., Jokisch, D., Gizewski, E. R., Forsting, M. & Daum, I. The neural coding of expected and unexpected monetary performance outcomes: Dissociations between active and observational learning. Behav. Brain Res. 227, 241–251 (2012).

42. Brewe, E. Modeling theory applied: Modeling Instruction in introductory physics. Am. J. Phys. 76, 1155–1160 (2008).

43. Huang, C.-C., Rolls, E. T., Feng, J. & Lin, C.-P. An extended Human Connectome Project multimodal parcellation atlas of the human cortex and subcortical areas. Brain Struct. Funct. 227, 763–778 (2022).

44. Glasser, M. F. et al. The Human Connectome Project’s neuroimaging approach. Nat. Neurosci. 19, 1175–1187 (2016).

45. Ji, J. L. et al. Mapping the human brain’s cortical-subcortical functional network organization. NeuroImage 185, 35–57 (2019).

46. Kong, R. Q. et al. A network correspondence toolbox for quantitative evaluation of novel neuroimaging results. Preprint at 10.1101/2024.06.17.599426 (2024).

47. Alexander-Bloch, A. F. et al. On testing for spatial correspondence between maps of human brain structure and function. NeuroImage 178, 540–551 (2018).

48. Bartley, J. E. et al. Brain activity links performance in science reasoning with conceptual approach. Npj Sci. Learn. 4, 20 (2019).

49. Singer, S.R, Nielsen, N., & Schweingruber, H. A. Discipline-Based Education Research: Understanding and Improving Learning in Undergraduate Science and Engineering. 13362 (National Academies Press, Washington, D.C., 2012). doi:10.17226/13362.

50. Shen, W. et al. Visual network alterations in brain functional connectivity in chronic low back pain: A resting state functional connectivity and machine learning study. NeuroImage Clin. 22, 101775 (2019).

51. Vossel, S., Geng, J. J. & Fink, G. R. Dorsal and Ventral Attention Systems: Distinct Neural Circuits but Collaborative Roles. The Neuroscientist 20, 150–159 (2014).

52. Mason, R. A. & Just, M. A. Neural Representations of Physics Concepts. Psychol. Sci. 27, 904–913 (2016).

53. Cohen, M. A., Horowitz, T. S. & Wolfe, J. M. Auditory recognition memory is inferior to visual recognition memory. Proc. Natl. Acad. Sci. 106, 6008–6010 (2009).

54. Pakan, J. M., Francioni, V. & Rochefort, N. L. Action and learning shape the activity of neuronal circuits in the visual cortex. Curr. Opin. Neurobiol. 52, 88–97 (2018).

55. Biswal, B., Zerrin Yetkin, F., Haughton, V. M. & Hyde, J. S. Functional connectivity in the motor cortex of resting human brain using echo-planar mri. Magn. Reson. Med. 34, 537–541 (1995).

56. Mackey, A. P., Miller Singley, A. T. & Bunge, S. A. Intensive Reasoning Training Alters Patterns of Brain Connectivity at Rest. J. Neurosci. 33, 4796–4803 (2013).

57. Mason, R. A. & Just, M. A. Physics instruction induces changes in neural knowledge representation during successive stages of learning. NeuroImage 111, 36–48 (2015).

58. Freedman, L., Zivan, M., Farah, R. & Horowitz-Kraus, T. Greater functional connectivity within the cingulo-opercular and ventral attention networks is related to better fluent reading: A resting-state functional connectivity study. NeuroImage Clin. 26, 102214 (2020).

59. Pearson, J. M., Heilbronner, S. R., Barack, D. L., Hayden, B. Y. & Platt, M. L. Posterior cingulate cortex: adapting behavior to a changing world. Trends Cogn. Sci. 15, 143–151 (2011).

60. Hill, N. M. & Schneider, W. Brain Changes in the Development of Expertise: Neuroanatomical and Neurophysiological Evidence about Skill-Based Adaptations. in The Cambridge Handbook of Expertise and Expert Performance (eds. Ericsson, K. A., Charness, N., Feltovich, P. J. & Hoffman, R. R.) 653–682 (Cambridge University Press, 2006). doi:10.1017/CBO9780511816796.037.

61. Jeon, H.-A. & Friederici, A. D. What Does “Being an Expert” Mean to the Brain? Functional Specificity and Connectivity in Expertise. Cereb. Cortex cercor;bhw329v1 (2016) doi:10.1093/cercor/bhw329.

62. Bjørndal, J. R., Beck, M. M., Jespersen, L., Christiansen, L. & Lundbye-Jensen, J. Hebbian priming of human motor learning. Nat. Commun. 15, 5126 (2024).

63. Cole, N., Harvey, M., Myers-Joseph, D., Gilra, A. & Khan, A. G. Prediction-error signals in anterior cingulate cortex drive task-switching. Nat. Commun. 15, 7088 (2024).

64. Allaire-Duquette, G. et al. An fMRI study of scientists with a Ph.D. in physics confronted with naive ideas in science. Npj Sci. Learn. 6, 11 (2021).

65. Fregni, S., Wolfensteller, U. & Ruge, H. Initial learning in the brain: From rules to action. Imaging Neurosci. 2, 1–34 (2024).

66. Leech, R. & Sharp, D. J. The role of the posterior cingulate cortex in cognition and disease. Brain 137, 12–32 (2014).

67. Trumpp, N. M., Ulrich, M. & Kiefer, M. Experiential grounding of abstract concepts: Processing of abstract mental state concepts engages brain regions involved in mentalizing, automatic speech, and lip movements. NeuroImage 288, 120539 (2024).

68. Abe, H. & Lee, D. Distributed Coding of Actual and Hypothetical Outcomes in the Orbital and Dorsolateral Prefrontal Cortex. Neuron 70, 731–741 (2011).

69. Mars, R. B. & Grol, M. J. Dorsolateral Prefrontal Cortex, Working Memory, and Prospective Coding for Action. J. Neurosci. 27, 1801–1802 (2007).

70. Wang, B. A., Veismann, M., Banerjee, A. & Pleger, B. Human orbitofrontal cortex signals decision outcomes to sensory cortex during behavioral adaptations. Nat. Commun. 14, 3552 (2023).

71. Jung-Beeman, M. et al. Neural Activity When People Solve Verbal Problems with Insight. PLoS Biol. 2, e97 (2004).

72. Qiu, J. et al. The neural basis of insight problem solving: An event-related potential study. Brain Cogn. 68, 100–106 (2008).

73. Federico, G. et al. The cortical thickness of the area PF of the left inferior parietal cortex mediates technical-reasoning skills. Sci. Rep. 12, 11840 (2022).

74. Ardila, A., Bernal, B. & Rosselli, M. Participation of the insula in language revisited: A meta-analytic connectivity study. J. Neurolinguistics 29, 31–41 (2014).

75. Chen, T. et al. Role of the anterior insular cortex in integrative causal signaling during multisensory auditory–visual attention. Eur. J. Neurosci. 41, 264–274 (2015).

76. Hamilton, L. S., Oganian, Y., Hall, J. & Chang, E. F. Parallel and distributed encoding of speech across human auditory cortex. Cell 184, 4626–4639.e13 (2021).

77. Vaden, K. I., Teubner-Rhodes, S., Ahlstrom, J. B., Dubno, J. R. & Eckert, M. A. Evidence for cortical adjustments to perceptual decision criteria during word recognition in noise. NeuroImage 253, 119042 (2022).

78. Khasawneh, E., Hodge-Zickerman, A., York, C. S., Smith, T. J. & Mayall, H. Examining the effect of inquiry-based learning versus traditional lecture-based learning on students’ achievement in college algebra. Int. Electron. J. Math. Educ. 18, em0724 (2023).

79. Sliwinska, M. W., James, A. & Devlin, J. T. Inferior Parietal Lobule Contributions to Visual Word Recognition. J. Cogn. Neurosci. 27, 593–604 (2015).

80. Koenigs, M., Barbey, A. K., Postle, B. R. & Grafman, J. Superior Parietal Cortex Is Critical for the Manipulation of Information in Working Memory. J. Neurosci. 29, 14980–14986 (2009).

81. Hosoda, C. et al. The structure of the superior and inferior parietal lobes predicts inter-individual suitability for virtual reality. Sci. Rep. 11, 23688 (2021).

82. Ishkhanyan, B. et al. Anterior and Posterior Left Inferior Frontal Gyrus Contribute to the Implementation of Grammatical Determiners During Language Production. Front. Psychol. 11, 685 (2020).

83. Bird, C. M. The role of the hippocampus in recognition memory. Cortex 93, 155–165 (2017).

84. Pergola, G., Ranft, A., Mathias, K. & Suchan, B. The role of the thalamic nuclei in recognition memory accompanied by recall during encoding and retrieval: An fMRI study. NeuroImage 74, 195–208 (2013).

85. Wolpert, D. M., Goodbody, S. J. & Husain, M. Maintaining internal representations: the role of the human superior parietal lobe. Nat. Neurosci. 1, 529–533 (1998).

86. Lydon-Staley, D. M., Ciric, R., Satterthwaite, T. D. & Bassett, D. S. Evaluation of confound regression strategies for the mitigation of micromovement artifact in studies of dynamic resting-state functional connectivity and multilayer network modularity. Netw. Neurosci. 3, 427–454 (2019).

87. Li, L., Zheng, Q., Xue, Y., Bai, M. & Mu, Y. Coactivation pattern analysis reveals altered whole-brain functional transient dynamics in autism spectrum disorder. Eur. Child Adolesc. Psychiatry (2024) doi:10.1007/s00787-024-02474-y.

88. Weber, S. et al. Transient resting-state salience-limbic co-activation patterns in functional neurological disorders. NeuroImage Clin. 41, 103583 (2024).

89. Zhang, R. et al. Disrupted brain state dynamics in opioid and alcohol use disorder: attenuation by nicotine use. Neuropsychopharmacology 49, 876–884 (2024).

90. Hestenes, D., Wells, M. & Swackhamer, G. Force concept inventory. Phys. Teach. 30, 141–158 (1992).

91. Lasry, N., Rosenfield, S., Dedic, H., Dahan, A. & Reshef, O. The puzzling reliability of the Force Concept Inventory. Am. J. Phys. 79, 909–912 (2011).

92. Von Korff, J., et al. Secondary analysis of teaching methods in introductory physics: A 50 k-student study. Am. J. Phys. 84, 969–974 (2016).

93. Elman, J. A., Klostermann, E. C., Marian, D. E., Verstaen, A. & Shimamura, A. P. Neural correlates of metacognitive monitoring during episodic and semantic retrieval. Cogn. Affect. Behav. Neurosci. 12, 599–609 (2012).

94. Avants, B., Epstein, C., Grossman, M. & Gee, J. Symmetric diffeomorphic image registration with cross-correlation: Evaluating automated labeling of elderly and neurodegenerative brain. Med. Image Anal. 12, 26–41 (2008).

95. Tustison, N. J. et al. N4ITK: Improved N3 Bias Correction. IEEE Trans. Med. Imaging 29, 1310–1320 (2010).

96. Esteban, O. et al. Analysis of task-based functional MRI data preprocessed with fMRIPrep. Nat. Protoc. 15, 2186–2202 (2020).

97. Reuter, M., Rosas, H. D. & Fischl, B. Highly accurate inverse consistent registration: A robust approach. NeuroImage 53, 1181–1196 (2010).

98. Gorgolewski, K. et al. Nipype: A Flexible, Lightweight and Extensible Neuroimaging Data Processing Framework in Python. Front. Neuroinformatics 5, (2011).

99. Dale, A. M., Fischl, B. & Sereno, M. I. Cortical Surface-Based Analysis. NeuroImage 9, 179–194 (1999).

100. Zhang, Y., Brady, M. & Smith, S. Segmentation of brain MR images through a hidden Markov random field model and the expectation-maximization algorithm. IEEE Trans. Med. Imaging 20, 45–57 (2001).

101. Greve, D. N. & Fischl, B. Accurate and robust brain image alignment using boundary-based registration. NeuroImage 48, 63–72 (2009).

102. Behzadi, Y., Restom, K., Liau, J. & Liu, T. T. A component based noise correction method (CompCor) for BOLD and perfusion based fMRI. NeuroImage 37, 90–101 (2007).

103. Satterthwaite, T. D. et al. An improved framework for confound regression and filtering for control of motion artifact in the preprocessing of resting-state functional connectivity data. NeuroImage 64, 240–256 (2013).

104. Esteban, O. et al. MRIQC: Advancing the automatic prediction of image quality in MRI from unseen sites. PLOS ONE 12, e0184661 (2017).

105. Atkinson, D., Hill, D. L. G., Stoyle, P. N. R., Summers, P. E. & Keevil, S. F. Automatic correction of motion artifacts in magnetic resonance images using an entropy focus criterion. IEEE Trans. Med. Imaging 16, 903–910 (1997).

106. Buxton, R. B., Uludağ, K., Dubowitz, D. J. & Liu, T. T. Modeling the hemodynamic response to brain activation. NeuroImage 23, S220–S233 (2004).

107. Wittkuhn, L. & Schuck, N. W. Dynamics of fMRI patterns reflect sub-second activation sequences and reveal replay in human visual cortex. Nat. Commun. 12, 1795 (2021).

108. Thorndike, R. L. Who belongs in the family? Psychometrika 18, 267–276 (1953).

109. Ingwersen, T. et al. Functional MRI brain state occupancy in the presence of cerebral small vessel disease **—** a pre-registered replication analysis of the Hamburg City Health Study. Imaging Neurosci. 2, 1–17 (2024).

110. Zhang, R. et al. Sleep Deprivation Effects on Brain State Dynamics Are Associated With Dopamine D2 Receptor Availability Via Network Control Theory. Biol. Psychiatry S0006322324015087 (2024) doi:10.1016/j.biopsych.2024.08.001.

111. Kupis, L. et al. Evoked and intrinsic brain network dynamics in children with autism spectrum disorder. NeuroImage Clin. 28, 102396 (2020).

112. Bates, D., Mächler, M., Bolker, B. & Walker, S. Fitting Linear Mixed-Effects Models Using **lme4**. J. Stat. Softw. 67, (2015).

113. Russell, Lenth. _emmeans: Estimated Marginal Means, aka Least-Squares Means_. (2024).

114. Smith, D. neurocaps. Zenodo 10.5281/ZENODO.14886003 (2025).

