## Supplemental Information for "Dynamic reconfiguration of brain coactivation states associated with active and lecture-based learning of university physics"

This document includes:

- Supplemental Methods
- Supplemental Results
- Supplemental Table 1
- Supplemental Table 2
- Supplemental Figure 1
- Supplemental Figure 2
- Supplemental References

### SUPPLEMENTAL METHODS

#### Neuroimaging Preprocessing

All functional magnetic resonance imaging (fMRI) data used in this study were preprocessed using `fMRIPrep` 1.5.0, a BIDS-compliant Python package which automatically provides a high-quality, reproducible preprocessing workflow with minimal user intervention[1].

##### Anatomical Data Preprocessing

T1-weighted images were corrected for intensity non-uniformity (INU) with `N4FieldCorrection`[2], distributed with `ANTs` 2.2.0 (RRID:SCR_004757)[3]. The T1w-reference was then skull-stripped with a `Nipype` implementation of the `antsBrainExtraction.sh` workflow (from `ANTs`), using `OASIS30ANTs` as target template. Brain tissue segmentation of the cerebrospinal fluid (CSF), white-matter (WM) and gray-matter (GM) was performed on the brain-extracted T1w using `fast` (`FSL` 5.0.9, RRIDLSCR_002823)[4]. A T1w-reference map was computed after registration of 2 T1w images (after INU-correction) using `mri_robust_template` (`FreeSurfer` 6.0.1)[5]. Brain surfaces were reconstructed using `recon-all` (`FreeSurfer` 6.0.1, RRID:SCR_001847)[6], and the brain mask estimated previously was refined with a custom variation of the method to reconcile `ANTs`-derived and `FreeSurfer`-derived segmentations of the cortical gray-matter of `Mindboggle` (RRID:SCR_002438)[7]. Volume-based spatial normalization to one standard space (MNI152NLin2009cAsym) was performed through nonlinear registration with `antsRegistration` (`ANTs` 2.2.0), using brain-extracted version of both T1w reference and the T1w template. The following template was selected for spatial normalization: ICBM 152 Nonlinear Asymmetrical template version 2009c {RRID:SCR_008796: TemplateFlow ID: MNI152NLin2009cAsym][8].

##### Functional Data Preprocessing

Each participant's dataset contained 1-3 runs of task-based functional magnetic resonance imaging (fMRI) data. For each BOLD run found per subject (across all tasks and sessions), the following preprocessing was performed. First, a reference volume and its skull-stripped version were generated using a custom methodology of `fMRIPrep`. A deformation field to correct for susceptibility distortions was estimated based on `fMRIPrep's` fieldmap-*less* approach. The deformation field is that resulting from co-registering the BOLD reference to the same-subject T1w-reference with its intensity inverted[9,10]. Registration is performed with antsRegistration (`ANTs` 2.2.0), and the process regularized by constraining deformation to be nonzero only along the phase-encoding direction, and modulated with an average fieldmap template[11]. Based on the estimated susceptibility distortion, an unwarped BOLD reference was calculated for a more accurate co-registration with the anatomical reference. The BOLD reference was then co-registered to the T1w reference using `bbregister` (`FreeSurfer`) which implements boundary-based registration[12]. Co-registration was configured with six degrees of freedom. Head-motion parameters with respect to the BOLD reference (transformation matrices, and six corresponding rotation and translation parameters) are estimated before any spatiotemporal filtering using `mcflirt` (`FSL` 5.0.9)[13]. The BOLD time-series were resampled to surfaces on the following spaces: *fsaverage5*. The BOLD time-series (including slice-timing correction when applied) were resampled onto their original, native space by applying a single, composite transform to correct for head-motion and susceptibility distortions. These resampled BOLD time-series will be referred to as preprocessed BOLD in original space, or just preprocessed BOLD. The BOLD time-series were resampled into standard space, generating a preprocessed BOLD run in *MNI152NLin2009cAsym* space. First, a reference volume and its skull-stripped version were generated using a custom methodology of fMRIPrep. Several confounding time-series were calculated based on the preprocessed BOLD: framewise displacement (FD), DVARS and three region-wise global signals. FD and DVARS are calculated for each functional run, both using their implementations in `Nipype` (following the definitions by Power et al[14]). The three global signals are extracted within the CSF, the WM, and the whole-brain masks. Additionally, a set of physiological regressors were extracted to allow for component-based noise correction (CompCor)[15]. Principal components are estimated after high-pass filtering the preprocessed BOLD time-series (using a discrete cosine filter with 128s cut-off) for the two CompCor variants: temporal (tCompCor) and anatomical (aCompCor). tCompCor components are then calculated from the top 5% variable voxels within a mask covering the subcortical regions. This subcortical mask is obtained by heavily eroding the brain mask, which ensures it does not include cortical GM regions. For aCompCor, components are calculated within the intersection of the aforementioned mask and the union of CSF and WM masks calculated in T1w space, after their projection to the native space of each functional run (using the inverse BOLD-to-T1w transformation). Components are also calculated separately within the WM and CSF masks. For each CompCor decomposition, the *k* components with the largest singular values are retained, such that the retained components' time series are sufficient to explain 50 percent of variance across the nuisance mask (CSF, WM, combined, or temporal). The remaining components are dropped from consideration. The head-motion estimates calculated in the correction step were also placed within the corresponding confounds file. The confound time series derived from head motion estimates and global signals were expanded with the inclusion of temporal derivatives and quadratic terms for each[16]. Frames that exceeded a threshold of 0.5 mm FD or 1.5 standardised DVARS were annotated as motion outliers. All resamplings can be performed with a single interpolation step by composing all the pertinent transformations (i.e. head-motion transform matrices, susceptibility distortion correction when available, and co-registrations to anatomical and output spaces). Gridded (volumetric) resamplings were performed using `antsApplyTransforms` (`ANTs`), configured with Lanczos interpolation to minimize the smoothing effects of other kernels[17]. Non-gridded (surface) resamplings were performed using `mri_vol2surf` (`FreesSurfer`).

#### SUPPLEMENTAL RESULTS

Prior to the CAPs analysis, functional images were excluded from the time series extraction process if MRIQC reports indicated that the mean framewise displacement, ghost-to-signal ratio, or entropy focus criterion exceeded the 99th percentile, or if the signal-to-noise ratio was below the 1st percentile; thus, indicating that these runs were outliers. Participants were also excluded if they did not possess at least one functional run for both pre-instruction and post-instruction resting state and task (FCI & PK) data. Based on these criteria, the sample size was reduced from 121 to 91 participants. Consequently, **Supplemental Table 1** presents the demographic information for the active learning (modeling-instruction) and lecture-based instruction groups after applying these exclusion criteria. As stated in the main manuscript, no significant differences between any of the demographic variables were noted between the active learning class or the lecture-based instruction class.

*Table S1.* **Participant Demographic Information After Exclusion Criteria.**

| | Modeling Instruction (*n*=46) | | Lecture-Based Instruction (*n*=44) | |
|---|---|---|---|---|
| | N | Percentage | N | Percentage |
| **Gender** | | | | |
| Male | 25 | 46 | 23 | 48 |
| Female | 21 | 54 | 21 | 52 |
| **Ethnicity** | | | | |
| Hispanic | 34 | 74 | 31 | 70 |
| Non-Hispanic | 12 | 26 | 13 | 30 |
| **Household Income** | | | | |
| < $15,000 | 8 | 17 | 9 | 20 |
| $15,000 - $34,999 | 10 | 22 | 10 | 23 |
| $35,000 - $49,999 | 5 | 11 | 6 | 14 |
| $50,000 - $74,999 | 12 | 26 | 5 | 11 |
| $75,000 - $99,999 | 9 | 20 | 6 | 14 |
| >$100,000 | 2 | 4 | 8 | 18 |
| **Years Enrolled** | | | | |
| Freshman | 3 | 6 | 5 | 11 |
| Sophomore | 24 | 52 | 20 | 45 |
| Junior | 14 | 20 | 12 | 27 |
| Senior | 5 | 11 | 7 | 16 |
| | **Grand Mean** | **Std. Dev.** | **Grand Mean** | **Std. Dev.** |
| **Age** | 19.96 | 1.76 | 19.89 | 1.50 |
| **GPA** | 3.31 | 0.62 | 3.25 | 0.41 |

*Note*. The "N" column represents the sample size of the group, and the "Percentage" column represents the percentage of participants in that group for each categorical variable (i.e., Gender, Ethnicity, Household Income, and Years Enrolled).

We conducted a comprehensive analysis of the demographic characteristics of students enrolled in the active learning classrooms (n=61) versus those in the lecture-based classrooms (n=60). The variables examined included age, sex, ethnicity, household income, grade point average (GPA), and academic standing at FIU (freshman, sophomore, junior, or senior). As detailed in **Supplementary Table 2**, statistical tests revealed no significant differences between the two groups across these demographics. Additionally, we focused on a subset of participants who completed at least one functional run for all tasks and sessions. This subsample comprised 46 students from the active learning classrooms and 44 from the lecture-based classrooms. Consistent with the broader sample, no significant demographic differences were observed between these groups (**Supplementary Table 2**)

**Table S2. Statistics for Demographic Information.**

| | Full Sample (n=121) | | Subsample (n=90) | |
|---|---|---|---|---|
| | Statistic | P-value | Statistic | P-value |
| **Sex** | $X^2(1) = 1.389$ | 0.239 | $X^2(1) = 0.174$ | 0.677 |
| **Ethnicity** | $X^2(1) = 0.00$ | 1.00 | $X^2(1) = 0.017$ | 0.896 |
| **Income** | $X^2(5) = 6.646$ | 0.248 | $X^2(5) = 7.191$ | 0.207 |
| **Years Enrolled** | $X^2(3) = 0.294$ | 0.961 | $X^2(3) = 1.307$ | 0.728 |
| **Pre-Instruction Age** | $t(116.84) = 0.362$ | 0.719 | $t(86.194) = 0.213$ | 0.832 |
| **Post-Instruction Age** | $t(117.73) = 0.301$ | 0.764 | $t(87.223) = 0.254$ | 0.800 |
| **Pre-Instruction GPA** | $t(95.319) = -0.068$ | 0.945 | $t(84.744) = 0.851$ | 0.397 |
| **Post-Instruction GPA** | $t(97.506) = 0.086$ | 0.932 | $t(73.558) = 0.266$ | 0.791 |

Once time series extraction and spatial dimensionality reduction using the HCPex parcellation were completed, each participant's functional runs from pre-instruction and post-instruction FCI, PK, and resting-state data were concatenated into a single matrix (Matrix rows: $N_{participants}*N_{TRs}$ ; Matrix columns: $N_{ROIs}$). This matrix was then subjected to k-means clustering with k values ranging from 2 to 20. The optimal value of $k$ was identified using the elbow method, which determines the point where inertia (the total within-cluster sum of squares) stops decreasing exponentially, indicating minimal information gain from higher values of $k$. As depicted in **Supplemental Figure 1**, the optimal $k$ value was found to be 7. Considering that the elbow method can be subjective due to its reliance on visual inspection, the `neurocaps` package utilizes the algorithm from `kneed`, a Python package, to select the elbow. This algorithm approximates the elbow of the plot by identifying the maximum and minimum coordinates, rotating the convex curve so that a horizontal line can be drawn between these coordinates, and then identifying the local maximum [18].

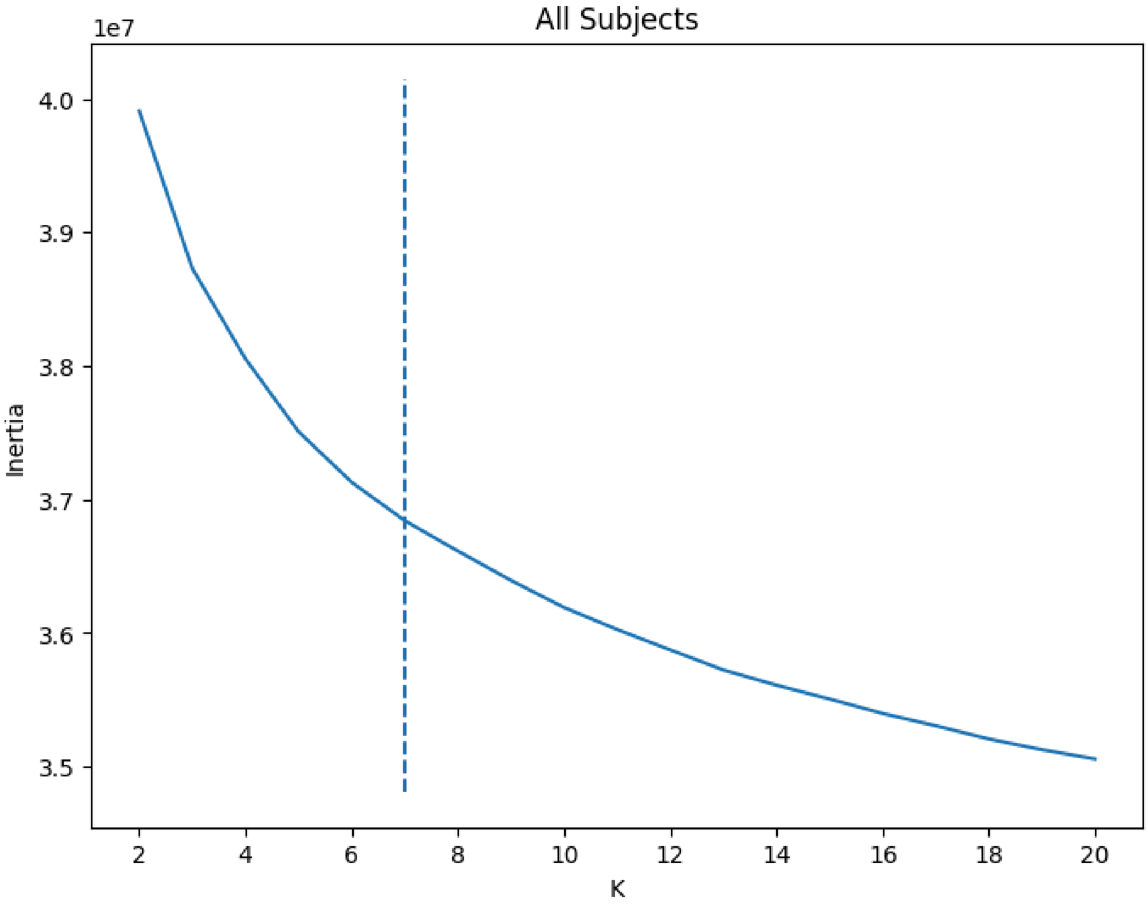


***Fig. S1. Inertia Values.*** A convex curve was constructed by plotting the inertia values for each *k* value. The dashed line indicates the optimal *k* value as identified by the kneed algorithm.

In this study, three reproducible CAP metrics[19] were calculated: counts, temporal fraction, and persistence. These metrics may be significantly influenced by the number of frames/time repetitions (TRs) each participant has. Any observed results for certain tasks, such as the FCI task - a self-paced task with a maximum duration of 45 seconds per block - could be affected by differences in the total number of frames per condition. Additionally, counts for the PK task and resting-state conditions may be influenced by scrubbing, which removes frames affected by excessive head motion, thereby compounding the issues observed in the FCI task. To mitigate these potential confounds, all analyses for the main effect of time, the main effect of class, and their interaction included the total number of frames per participant as an additional covariate. This adjustment ensures that differences in counts are attributable to time, class, or their interaction rather than variations in the number of frames. Furthermore, t-tests (paired t-test for the time comparison and independent t-test for the class comparison) were conducted to evaluate whether the total number of frames differed significantly across classes and time points for each task. The results of these tests are detailed in **Supplemental Table 3.**

**Table S3. Descriptive Information of Total Frames/TRs.**

| Paradigm | Comparison | Group | N | Mean | Std. Dev. | T-Statistic | P |
|---|---|---|---|---|---|---|---|
| FCI | Time | Pre-Instruction | 90 | 114.76 | 19.97 | 3.842 | **<0.001*** ** |
| | | Post-Instruction | 90 | 104.34 | 21.22 | | |
| | Class (Pre-Instruction) | Modeling | 46 | 114.65 | 21.42 | 0.050 | 0.960 |
| | | Lecture | 44 | 114.86 | 18.32 | | |
| | Class (Post-Instruction) | Modeling | 46 | 104.17 | 20.37 | 0.078 | 0.938 |
| | | Lecture | 44 | 104.52 | 22.06 | | |
| PK | Time | Pre-Instruction | 90 | 88.30 | 6.81 | 0.365 | 0.491 |
| | | Post-Instruction | 90 | 87.43 | 9.47 | | |
| | Class (Pre-Instruction) | Modeling | 46 | 88.26 | 6.90 | 0.055 | 0.956 |
| | | Lecture | 44 | 88.34 | 6.70 | | |
| | Class (Post-Instruction) | Modeling | 46 | 87.52 | 9.17 | 0.090 | 0.929 |
| | | Lecture | 44 | 87.34 | 9.77 | | |
| Resting-State | Time | Pre-Instruction | 90 | 353.19 | 10.72 | 1.380 | 0.171 |
| | | Post-Instruction | 90 | 350.19 | 22.20 | | |
| | Class (Pre-Instruction) | Modeling | 46 | 356.04 | 6.07 | 2.65 | **0.009*** ** |
| | | Lecture | 44 | 350.20 | 13.38 | | |
| | Class (Post-Instruction) | Modeling | 46 | 354.06 | 18.11 | 1.702 | 0.092 |
| | | Lecture | 44 | 346.14 | 25.16 | | |

*Note.* P-values accompanied by an asterisk denotes values less than 0.05, signifying that the groups differ in their total number of frames.

Each metric (counts, temporal fraction, and persistence) computed for each of the seven CAPs was analyzed to assess the main effect of time (**Supplemental Table 4**), the main effect of class (**Supplemental Table 5**), and their interaction effects (**Supplemental Table 6**). The main effect of time was evaluated by contrasting post-instruction with pre-instruction data. The main effect of class was assessed by contrasting active learning with lecture-based instruction. For the interaction effect, the lecture-based instruction group served as the reference group. Additionally, all contrasts were corrected for multiple comparisons using the Benjamini-Hochberg procedure.

**Table S4. Main Effect of Time.**

| Metrics | Task | | |
|---|---|---|---|
| | FCI | PK | Resting State |
| **CAP-1** | | | |
| Counts | Δ = -0.399, (p = 0.151) | Δ = -0.330, (p = 0.171) | **Δ = -1.942***, (p < 0.001)** |
| Temporal Fraction | Δ = -0.010, (p = 0.079) | Δ = -0.007, (p = 0.314) | **Δ = -0.019***, (p < 0.001)** |
| Persistence | Δ = -0.085, (p = 0.565) | Δ = 0.037, (p = 0.778) | **Δ = -0.202, (p = 0.036)** |
| **CAP-2** | | | |
| Counts | Δ = -0.435, (p = 0.130) | Δ = -0.157, (p = 0.507) | Δ = -0.307, (p = 0.486) |
| Temporal Fraction | Δ = -0.007, (p = 0.226) | Δ = -0.004, (p = 0.542) | Δ = 0.001, (p = 0.820) |
| Persistence | Δ = 0.006, (p = 0.964) | Δ = -0.034, (p = 0.805) | Δ = 0.111, (p = 0.243) |
| **CAP-3** | | | |
| Counts | Δ = -0.013, (p = 0.964) | Δ = 0.501, (p = 0.094) | Δ = 0.673, (p = 0.201) |
| Temporal Fraction | Δ = -0.008, (p = 0.146) | Δ = 0.015, (p = 0.083) | Δ = 0.006, (p = 0.198) |
| Persistence | Δ = -0.124, (p = 0.149) | Δ = 0.201, (p = 0.162) | Δ = 0.041, (p = 0.656) |
| **CAP-4** | | | |
| Counts | Δ = 0.007, (p = 0.98) | Δ = -0.234, (p = 0.326) | **Δ = -1.087*, (p = 0.011)** |
| Temporal Fraction | Δ = 0.002, (p = 0.738) | Δ = -0.009, (p = 0.104) | **Δ = -0.012**, (p < 0.001)** |
| Persistence | Δ = 0.004, (p = 0.970) | Δ = -0.175, (p = 0.207) | **Δ = -0.210, (p = 0.025)** |
| **CAP-5** | | | |
| Counts | Δ = -0.150, (p = 0.642) | **Δ = 0.621, (p = 0.032)** | Δ = -0.174, (p = 0.728) |
| Temporal Fraction | Δ = -0.002, (p = 0.801) | **Δ = 0.019, (p = 0.010)** | Δ = 0.003, (p = 0.333) |
| Persistence | Δ = -0.001, (p = 0.990) | Δ = 0.157, (p = 0.221) | Δ = 0.117, (p = 0.113) |
| **CAP-6** | | | |
| Counts | Δ = 0.507, (p = 0.064) | Δ = -0.284, (p = 0.177) | Δ = 0.365, (p = 0.437) |
| Temporal Fraction | **Δ = 0.014, (p = 0.018)** | Δ = -0.005, (p = 0.438) | Δ = 0.008, (p = 0.057) |
| Persistence | Δ = 0.076, (p = 0.560) | Δ = -0.029, (p = 0.816) | Δ = 0.165, (p = 0.057) |
| **CAP-7** | | | |
| Counts | Δ = 0.496, (p = 0.101) | Δ = -0.234, (p = 0.364) | **Δ = 1.129*, (p = 0.019)** |
| Temporal Fraction | Δ = 0.011, (p = 0.082) | Δ = -0.009, (p = 0.172) | **Δ = 0.014**, (p = 0.001)** |
| Persistence | Δ = 0.042, (p = 0.724) | Δ = -0.005, (p = 0.972) | **Δ = 0.203, (p = 0.047)** |

*Note.* Delta (Δ) represents change from pre- to post-instruction. P-values in parentheses are uncorrected for multiple comparisons. Significant uncorrected p-values are bold. Asterisks indicate significance after Benjamini-Hochberg Correction: $*p_{BH} < 0.05$, $**p_{BH} < .01$, $***p_{BH} < 0.001$.

**Table S5. Main Effect of Instruction.**

| Metrics | Task | | |
|---|---|---|---|
| | FCI | PK | Resting State |
| **CAP-1** | | | |
| Counts | Δ = 0.231, (p = 0.391) | Δ = 0.100, (p = 0.694) | Δ = 0.470, (p = 0.394) |
| Temporal Fraction | Δ = 0.00, (p = 0.996) | Δ = 0.009, (p = 0.161) | Δ = 0.007, (p = 0.155) |
| Persistence | Δ = -0.132, (p = 0.362) | Δ = 0.226, (p = 0.130) | Δ = 0.146, (p = 0.132) |
| **CAP-2** | | | |
| Counts | Δ = -0.489, (p = 0.113) | Δ = -0.327, (p = 0.170) | Δ = -0.060, (p = 0.896) |
| Temporal Fraction | Δ = 0.00, (p = 0.994) | Δ = 0.002, (p = 0.706) | Δ = -0.003, (p = 0.413) |
| Persistence | **Δ = 0.267, (p = 0.046)** | **Δ = 0.346, (p = 0.014)** | Δ = -0.092, (p = 0.437) |
| **CAP-3** | | | |
| Counts | Δ = 0.229, (p = 0.432) | Δ = -0.22, (p = 0.494) | Δ = 0.064, (p = 0.914) |
| Temporal Fraction | Δ = -0.002, (p = 0.737) | Δ = 0.01, (p = 0.232) | Δ = -0.003, (p = 0.503) |
| Persistence | Δ = -0.100, (p = 0.334) | **Δ = 0.394, (p = 0.007)** | Δ = -0.092, (p = 0.347) |
| **CAP-4** | | | |
| Counts | Δ = 0.252, (p = 0.352) | Δ = -0.154, (p = 0.519) | Δ = 0.404, (p = 0.490) |
| Temporal Fraction | Δ = 0.003, (p = 0.653) | Δ = -0.003, (p = 0.635) | Δ = 0.004, (p = 0.314) |
| Persistence | Δ = -0.057, (p = 0.69) | Δ = 0.061, (p = 0.661) | Δ = 0.049, (p = 0.601) |
| **CAP-5** | | | |
| Counts | Δ = 0.049, (p = 0.878) | Δ = -0.329, (p = 0.256) | Δ = 0.381, (p = 0.474) |
| Temporal Fraction | Δ = -0.003, (p = 0.597) | Δ = -0.006, (p = 0.423) | Δ = 0.00, (p = 0.948) |
| Persistence | Δ = -0.073, (p = 0.479) | Δ = 0.033, (p = 0.827) | Δ = -0.078, (p = 0.430) |
| **CAP-6** | | | |
| Counts | Δ = -0.123, (p = 0.684) | Δ = -0.153, (p = 0.565) | Δ = 0.225, (p = 0.711) |
| Temporal Fraction | Δ = 0.003, (p = 0.596) | Δ = -0.005, (p = 0.498) | Δ = 0.003, (p = 0.572) |
| Persistence | Δ = 0.203, (p = 0.152) | Δ = -0.132, (p = 0.341) | Δ = 0.030, (p = 0.795) |
| **CAP-7** | | | |
| Counts | Δ = 0.292, (p = 0.318) | Δ = 0.023, (p = 0.933) | Δ = -0.811, (p = 0.118) |
| Temporal Fraction | Δ = -0.001, (p = 0.928) | Δ = -0.007, (p = 0.349) | Δ = -0.008, (p = 0.09) |
| Persistence | Δ = -0.133, (p = 0.318) | Δ = -0.208, (p = 0.161) | Δ = -0.078, (p = 0.478) |

*Note.* Delta (Δ) represents the contrast between the active learning and lecture-based instruction groups. P-values in parentheses are uncorrected for multiple comparisons. Significant uncorrected p-values are bold. Asterisks indicate significance after Benjamini-Hochberg Correction: $^{**}p_{BH} < 0.05$, $^{***}p_{BH} < 0.001$.

**Table S6. Interaction Effect.**

| Metrics | Task | | |
|---|---|---|---|
| | FCI | PK | Resting State |
| **CAP-1** | | | |
| Counts | Δ = 0.092, (p = 0.863) | Δ = -0.130, (p = 0.786) | Δ = -0.447, (p = 0.616) |
| Temporal Fraction | Δ = -0.006, (p = 0.569) | Δ = 0.006, (p = 0.628) | Δ = 0.013, (p = 0.087) |
| Persistence | Δ = -0.234, (p = 0.416) | Δ = 0.233, (p = 0.374) | **Δ = 0.494, (p = 0.010)** |
| **CAP-2** | | | |
| Counts | Δ = -0.976, (p = 0.079) | Δ = -0.298, (p = 0.530) | Δ = 0.325, (p = 0.711) |
| Temporal Fraction | Δ = -0.006, (p = 0.592) | Δ = -0.008, (p = 0.528) | Δ = 0.005, (p = 0.455) |
| Persistence | Δ = 0.404, (p = 0.128) | Δ = -0.255, (p = 0.356) | Δ = 0.103, (p = 0.586) |
| **CAP-3** | | | |
| Counts | Δ = -0.621, (p = 0.285) | Δ = 0.633, (p = 0.288) | Δ = 0.586, (p = 0.574) |
| Temporal Fraction | Δ = -0.005, (p = 0.669) | Δ = 0.015, (p = 0.355) | Δ = 0.006, (p = 0.485) |
| Persistence | Δ = 0.215, (p = 0.191) | Δ = 0.034, (p = 0.904) | Δ = 0.059, (p = 0.746) |
| **CAP-4** | | | |
| Counts | Δ = -0.762, (p = 0.159) | Δ = -0.336, (p = 0.478) | Δ = 0.612, (p = 0.466) |
| Temporal Fraction | Δ = 0.010, (p = 0.355) | Δ = -0.001, (p = 0.963) | Δ = -0.002, (p = 0.810) |
| Persistence | **Δ = 0.549, (p = 0.013)** | Δ = 0.185, (p = 0.504) | Δ = -0.208, (p = 0.261) |
| **CAP-5** | | | |
| Counts | Δ = -0.03, (p = 0.961) | Δ = 0.399, (p = 0.486) | Δ = 0.487, (p = 0.624) |
| Temporal Fraction | Δ = 0.007, (p = 0.593) | Δ = 0.008, (p = 0.560) | Δ = -0.002, (p = 0.762) |
| Persistence | Δ = 0.183, (p = 0.374) | Δ = -0.018, (p = 0.943) | Δ = -0.103, (p = 0.480) |
| **CAP-6** | | | |
| Counts | Δ = 0.198, (p = 0.705) | Δ = 0.15, (p = 0.721) | **Δ = -1.859, (p = 0.048)** |
| Temporal Fraction | Δ = 0.002, (p = 0.882) | Δ = -0.011, (p = 0.347) | **Δ = -0.016, (p = 0.038)** |
| Persistence | Δ = -0.175, (p = 0.489) | Δ = -0.433, (p = 0.087) | Δ = -0.159, (p = 0.350) |
| **CAP-7** | | | |
| Counts | Δ = -0.449, (p = 0.441) | **Δ = -1.124, (p = 0.031)** | Δ = -0.284, (p = 0.764) |
| Temporal Fraction | Δ = -0.002, (p = 0.898) | Δ = -0.011, (p = 0.408) | Δ = -0.004, (p = 0.621) |
| Persistence | Δ = 0.045, (p = 0.846) | Δ = 0.466, (p = 0.115) | Δ = -0.041, (p = 0.840) |

*Note.* Delta (Δ) represents the relative change of the active learning group relative to the reference group, the lecture-based instruction group. P-values in parentheses are uncorrected for multiple comparisons. Significant uncorrected p-values are bold. Asterisks indicate significance after Benjamini-Hochberg Correction: **$p_{BH} < 0.05$, *** $p_{BH} < 0.001$.

A fourth metric, transition probability, was also identified as a reproducible CAP metric by [19]. Transition probability computes the probability of transitioning from one state to another or remaining in the same state and is calculated as the number of transitions from State A to State B divided by the total transitions from State A ($\frac{N_{A\rightarrow B}}{N_A}$). This metric provides insight into which CAPs are more likely to transition from specific CAPs. Despite its informative value, using this metric would result in 441 additional statistical tests (49 elements multiplied by three contrasts - main effect of time, main effect of class, and interaction effect - conducted for Resting State, the FCI task, and the PK task). Although these statistical tests were not performed, transition probabilities were calculated for each participant. The average transition matrices for the FCI task, and the PK task, and resting state are shown for both the active learning and lecture-based instruction groups at pre-instruction and post-instruction sessions. **Supplemental Figure 2** is provided to give readers a foundational understanding of how CAP transitions occur for each class.

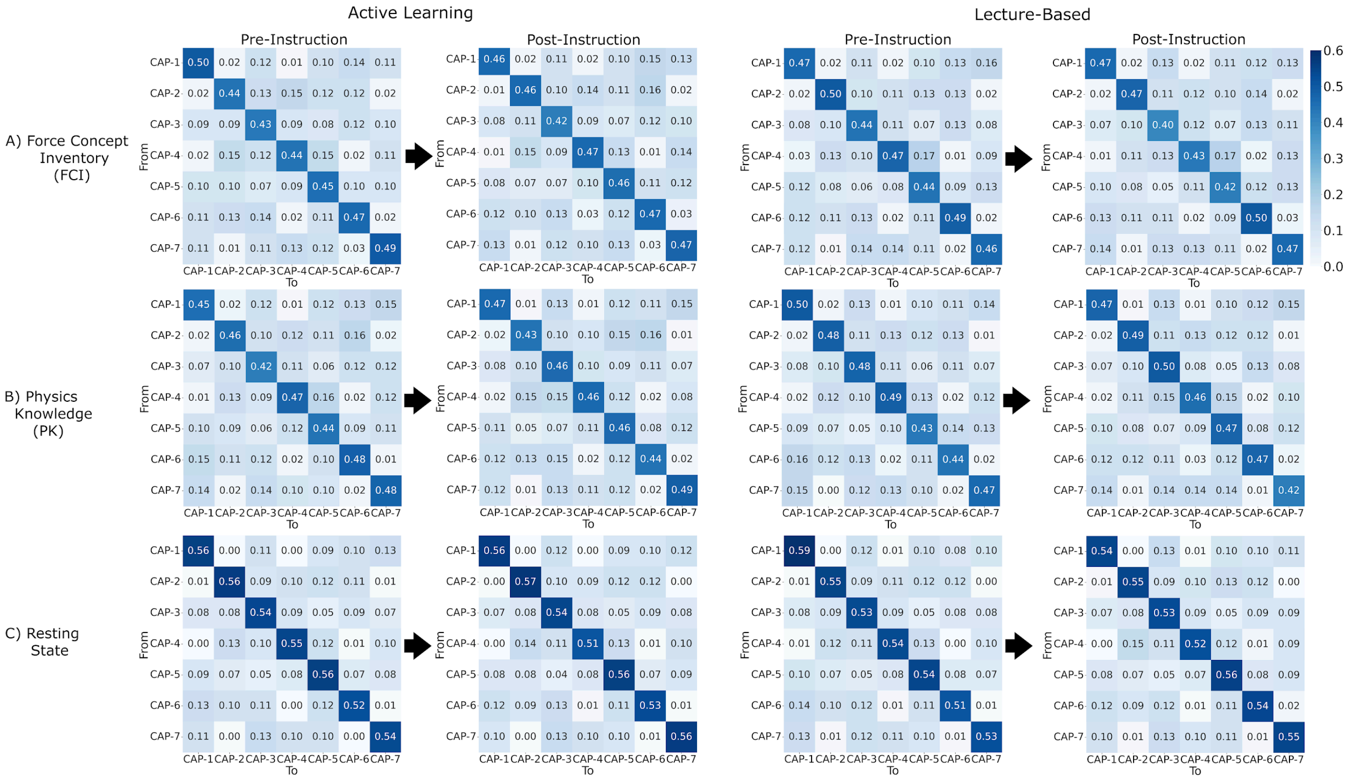


***Figure S2. Transition Probability.*** Each heatmap shows the averaged transition probability matrix for the corresponding session (pre-instruction vs. post-instruction), fMRI task (FCI, PK, & Resting State), and class type (active learning vs. lecture-based instruction). Colors range from light blue to dark blue, with darker hues indicating higher transition probabilities. The vertical columns represent the "from" states, while the horizontal rows indicate the "to" states. A higher-resolution image is available on the project OSF page.

#### Supplemental References


1. Esteban, O. *et al.* fMRIPrep: a robust preprocessing pipeline for functional MRI. *Nat. Methods* **16**, 111–116 (2019).
2. Tustison, N. J. *et al.* N4ITK: Improved N3 Bias Correction. *IEEE Trans. Med. Imaging* **29**, 1310–1320 (2010).
3. Avants, B., Epstein, C., Grossman, M. & Gee, J. Symmetric diffeomorphic image registration with cross-correlation: Evaluating automated labeling of elderly and neurodegenerative brain. *Med. Image Anal.* **12**, 26–41 (2008).
4. Zhang, Y., Brady, M. & Smith, S. Segmentation of brain MR images through a hidden Markov random field model and the expectation-maximization algorithm. *IEEE Trans. Med. Imaging* **20**, 45–57 (2001).
5. Reuter, M., Rosas, H. D. & Fischl, B. Highly accurate inverse consistent registration: A robust approach. *NeuroImage* **53**, 1181–1196 (2010).
6. Dale, A. M., Fischl, B. & Sereno, M. I. Cortical Surface-Based Analysis. *NeuroImage* **9**, 179–194 (1999).
7. Klein, A. *et al.* Mindboggling morphometry of human brains. *PLOS Comput. Biol.* **13**, e1005350 (2017).
8. Fonov, V. *et al.* Unbiased average age-appropriate atlases for pediatric studies. *NeuroImage* **54**, 313–327 (2011).
9. Huntenburg, J. M. Evaluating Nonlinear Coregistration of BOLD EPI and T1w Images. *Berl. Freie Univ.* (2014).
10. Wang, S. *et al.* Evaluation of Field Map and Nonlinear Registration Methods for Correction of Susceptibility Artifacts in Diffusion MRI. *Front. Neuroinformatics* **11**, (2017).
11. Treiber, J. M. *et al.* Characterization and Correction of Geometric Distortions in 814 Diffusion Weighted Images. *PLOS ONE* **11**, e0152472 (2016).
12. Greve, D. N. & Fischl, B. Accurate and robust brain image alignment using boundary-based registration. *NeuroImage* **48**, 63–72 (2009).
13. Jenkinson, M., Bannister, P., Brady, M. & Smith, S. Improved Optimization for the Robust and Accurate Linear Registration and Motion Correction of Brain Images. *NeuroImage* **17**, 825–841 (2002).
14. Power, J. D. *et al.* Methods to detect, characterize, and remove motion artifact in resting state fMRI. *NeuroImage* **84**, 320–341 (2014).
15. Behzadi, Y., Restom, K., Liau, J. & Liu, T. T. A component based noise correction method (CompCor) for BOLD and perfusion based fMRI. *NeuroImage* **37**, 90–101 (2007).
16. Satterthwaite, T. D. *et al.* An improved framework for confound regression and filtering for control of motion artifact in the preprocessing of resting-state functional connectivity data. *NeuroImage* **64**,

240–256 (2013).

17. Lanczos, C. Evaluation of Noisy Data. *J. Soc. Ind. Appl. Math. Ser. B Numer. Anal.* **1**, 76–85 (1964).
18. Satopaa, V., Albrecht, J., Irwin, D. & Raghavan, B. Finding a 'Kneedle' in a Haystack: Detecting Knee Points in System Behavior. in *2011 31st International Conference on Distributed Computing Systems Workshops* 166–171 (IEEE, Minneapolis, MN, USA, 2011). doi:10.1109/ICDCSW.2011.20.
19. Yang, H. *et al.* Reproducible coactivation patterns of functional brain networks reveal the aberrant dynamic state transition in schizophrenia. *NeuroImage* **237**, 118193 (2021).
